# ClonoScreen3D: a novel three-dimensional clonogenic screening platform for identification of radiosensitizers for glioblastoma

**DOI:** 10.1101/2023.10.04.560635

**Authors:** Mark R Jackson, Amanda R Richards, Abdul-Basit Ayoola Oladipupo, Sandeep K Chahal, Seamus Caragher, Anthony J Chalmers, Natividad Gomez-Roman

**Author notes:** NC3Rs project grant, Grant Ref: NC/P001335/1. Conflict of interest: The authors declare no potential conflicts of interest. Corresponding author: Natividad Gomez-Roman, Strathclyde Institute of Pharmacy and Biomedical Sciences, University of Strathclyde, 161 Cathedral Street, Glasgow G4 0RE.

## Abstract

**Purpose:** Glioblastoma (GBM) is a lethal brain tumour. Standard of care treatment comprising surgery, radiation and chemotherapy results in median survival rates of 12-15 months. Molecular targeted agents identified using conventional two-dimensional (2D) *in vitro* models of GBM have failed to improve outcome in patients, rendering such models inadequate for therapeutic target identification. We developed a 3D GBM *in vitro* model that recapitulates key GBM clinical features and responses to molecular therapies and investigated its utility for screening novel radiation-drug combinations using gold-standard clonogenic survival as readout.

**Results:** Patient-derived GBM cell lines were optimized for inclusion in a 96-well plate 3D clonogenic screening platform, ClonoScreen3D. Radiation responses of GBM cells in this system were highly reproducible and comparable to those observed in low-throughout 3D assays. The screen methodology provided quantification of candidate drug single agent activity (EC_50_) and the interaction between drug and radiation (radiation interaction ratio, RIR). The PARP inhibitors talazoparib, rucaparib and olaparib, each showed a significant interaction with radiation by ClonoScreen3D and were subsequently confirmed as true radiosensitizers by full clonogenic assay. Screening a panel of DNA damage response inhibitors revealed the expected propensity of these compounds to interact significantly with radiation (13/15 compounds). A second screen assessed a panel of compounds targeting pathways identified by transcriptomic analysis and demonstrated single agent activity and a previously unreported interaction with radiation of dinaciclib and cytarabine (RIR 1.28 and 1.90, respectively). These compounds were validated as radiosensitizers in full clonogenic assays (sensitizer enhancement ratio 1.47 and 1.35, respectively).

**Conclusions:** The ClonoScreen3D platform was demonstrated to be a robust method to screen for single agent and radiation-drug combination activity. Using gold-standard clonogenicity, this assay is a tool for identification of novel radiosensitizers. We anticipate this technology will accelerate identification of novel radiation-drug combinations with genuine translational value.

## Introduction

Glioblastoma (GBM) is the most common and most aggressive primary brain tumour [1]. Even with trimodal therapy comprising surgery, radiation and chemotherapy (temozolomide) [2], prognosis remains dismal owing to marked chemo– and radioresistance [3]. Multiple molecularly targeted agents exploiting pathways commonly dysregulated in GBM, including the receptor tyrosine kinase/Ras/phosphoinositide 3-kinase (PI3K) [3–5], αv integrins [6], p53 and retinoblastoma pathways, have shown therapeutic efficacy in preclinical models of GBM. However, all these agents have failed in the clinic, either alone or in combination with standard of care radiotherapy and/or chemotherapy [3, 5, 7, 8]. These results emphasise the need for improved experimental models that translate effectively to the clinic. We developed a novel three-dimensional (3D) *in vitro* GBM culture system that better reflects patient response to treatment [9] and has been extensively characterized in terms of cell morphology, mRNA and protein expression, and response to therapies including radiation, chemotherapy (temozolomide) and clinically-relevant molecular targeted agents (e.g. erlotinib, bevacizumab). Comparison of the 3D system with conventional 2D or neurosphere models has confirmed its superiority for drug discovery in GBM [10].

Radiotherapy is a central component of GBM treatment and while its efficacy in terms of overall survival has been proven in clinical trials [11], tumour recurrence occurs in nearly all patients. Radiation dose escalation has not improved clinical outcomes and most patients experience disabling neurocognitive toxicity. Enhancing the efficacy of radiotherapy will therefore require the use of radiosensitizing drugs that potentiate cytotoxicity in a tumour-specific manner. Radioresistance in GBM has been linked to a subpopulation of cells termed GBM stem-like cells, which display preferential activation of the DNA damage response (DDR) and increased DNA repair capacity [12, 13]. Pharmacological disruption of the DDR therefore offers an attractive strategy to overcome radioresistance and enable radiation therapy to eliminate this problematic population of tumour cells.

Identification of radiosensitizers *in vitro* is best achieved using the gold-standard clonogenic survival assay (CSA), but this is currently labour-intensive and time-consuming. Furthermore, difficulties in compressing 2D CSAs to small growth area formats (e.g. 96-well plate) due to colony size has presented a barrier [14, 15]. Nevertheless, in addition to non-clonogenic screening methods, efforts to establish clonogenic or pseudoclonogenic platforms have been reported [14–18] in addition to non-clonogenic screening methods. However, development of a medium or high-throughput screen to identify radiosensitizers using primary GBM cells cultured in 3D conditions has not previously been reported.

To overcome these obstacles, we modified our clinically-relevant 3D clonogenic system to 96-well plate format and observed that the high surface area of the 3D-Alvetex scaffold supported growth of sufficient numbers of colonies for large-scale compound screening. To identify novel, clinically exploitable targets for radiosensitization, we performed RNAseq analysis of GBM cells grown in 3D before and after radiation treatment. In addition to the expected DDR candidates, we identified novel potential targets in pathways including cell cycle progression, mitosis, and DNA synthesis. Our novel screening tool, the ClonoScreen3D platform, represents an improved experimental strategy for streamlining identification of novel radiosensitizers and has potential to transform the landscape of GBM therapy.

## Materials and Methods

### Cell culture and treatment

Patient-derived G7 and E2 GBM cells were obtained from Professor Colin Watts, as previously described [19]. Patient-derived GBML20 GBM cells were obtained Dr Dimitris Placantonakis (NYU). Cells were cultured as monolayers on Matrigel-coated plates (0.2347 mg/mL in Adv/DMEM) in cancer stem cell enriching serum-free medium comprising Advanced/DMEM/F12 medium (GIBCO) supplemented with 1% B27 and 0.5% N2 (Thermo Fisher Scientific), 4 μg/mL heparin, 10 ng/mL fibroblast growth factor 2 (bFGF, Merck/Sigma), 20 ng/mL epidermal growth factor (EGF, Sigma) and 1% L-glutamine. G7s cells were grown in suspension conditions as spheres for routine passage and were then grown as monolayers on Matrigel-coated plates for 4-7 passages prior to seeding on 3D-Alvetex plates. G7m cells were maintained as monolayers on Matrigel-coated plates. Despite originating from the same parental lines, these two models exhibit distinct features, including radiation responses. Cell lines were grown for a maximum of seven passages before inclusion in experiments at 37°C, with 5% CO_2_. All cells were routinely monitored for mycoplasma contamination.

For 3D-Alvetex (Reprocell) cultures, plates were pre-treated according to the manufacturer’s instructions (washed with 70% ethanol, followed by three washes with PBS). 3D-Alvetex scaffolds were coated with Matrigel (0.2347 mg/mL), using 50 μL/well for 96-well 3D-Alvetex plates or 0.5 ml/well for 12-well 3D-Alvetex plates.

### ClonoScreen3D clonogenic survival assay

Seeding densities were as follows: 180 cells/well for G7s and 150 cells/well for G7m cells. Eighteen hours after seeding, cells were treated with vehicle (DMSO) or inhibitor for 2 hours. Cells were irradiated (3 Gy) using an RS225 (XStrahl) X-ray cabinet, at 195 kV, 15 mA with a 0.5 mm copper filter, at a dose rate of 2.47 Gy/min, or sham-irradiated. Details of the inhibitors used are listed in Supplementary Table 1. Colonies were grown for 14 days at 37°C, 5% CO_2_, followed by incubation with thiazolyl blue tetrazolium bromide (MTT) for 4 hours at 37°C. Cells were fixed using 2% paraformaldehyde in PBS at room temperature for 15 min and washed with PBS.

### Image acquisition and processing

High resolution images of plates were acquired using a photographic set-up comprising a white transilluminator to optimise contrast (Voliamart A3 Tracing Board) with a camera downward copy stand carrying a digital camera (Nikon D5300+AF-P 18-55VR). Image capture was performed using digiCamControl software. Well segmentation was performed with ImageJ software. Colonies composed of >50 cells were counted either manually or in a semi-automated manner using the open-source software, OpenCFU (http://opencfu.sourceforge.net).

### Quantification of single agent activity and radiosensitizing potential

Colony counts were converted into surviving fraction (SF) using the plating efficiency of vehicle-treated cells for each radiation dose, thus correcting the values for the effect of ionizing radiation (IR) [20] [21, 22]. For determination of single agent activity, the mean SFs of sham irradiated replicates were calculated and modelled using a 4-parameter dose response curve, using the ‘drc’ package [22] in R (3.6.3) (https://www.R-project.org). Single agent activity was expressed in terms of EC_50_. For compounds that lacked single agent activity or did not conform to a classical dose response, EC_50_ was not determined.

Radiosensitizing potential was quantified by calculation of a novel parameter: the radiation interaction ratio (RIR). The linear interpolation area-under-the-curve (AUC) of SF against log_10_(drug concentration) was computed for individual biological replicates at each radiation dose, using the ‘MESS’ package [23]. The replicate AUCs were subjected to a ratio t-test from the ‘mratios’ package [24] with ratio under the null hypothesis ρ=1, to determine the statistical significance of the interaction with IR. P-values were adjusted for multiple comparison using the false discovery rate method. Thus, RIR was defined as

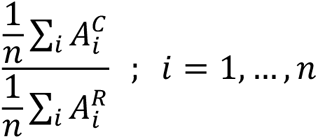

where *A^C^* is the AUC of the sham irradiated control and *A^R^* is the AUC of the irradiated sample. An example R script for computation of RIR can be found as supplementary material. Where data conformed to a dose response curve, the EC_50_ values of the sham and irradiated samples were additionally compared by t-test of coefficient ratio.

## 3D clonogenic survival assay for full radiation dose response

12-well plate 3D-Alvetex CSA were performed as previously described [22]. Briefly, seeding densities for all cell lines cells varied according to radiation dose: 300 cells/well for 0, 1, and 2 Gy; 500 cells/well for 3 Gy; 800 cells/well for 4 Gy and 1000 cells/well for 5 Gy. Eighteen hours after seeding, cells were treated with vehicle (DMSO) or inhibitor for 2 hours at 37°C (5% CO_2_) and subsequently sham-irradiated or irradiated at different radiation doses (1-5 Gy). Colonies were grown for 18-21 days at 37°C, 5% CO_2_, followed by incubation with thiazolyl blue tetrazolium bromide (MTT) for 4 hours at 37°C and fixed with 2% paraformaldehyde in PBS at room temperature for 15 min and washed with PBS. Plates were stored in PBS at 4°C, which was removed immediately prior to imaging and automated colony counting. Drug sensitizer enhancement ratios (SER) were calculated using mean inactivation doses determined from linear quadratic fits as described in [23].

### Gene expression analysis

Four days after plating cells in 3D conditions (3D-Alvetex), RNA was extracted with TRIzol reagent. RNAseq analysis was performed using the IlluminaNextSeq500 for a PolyA selection RNA library, with a paired-end sequencing model and 33M depth for triplicate experimental repeats of 2D and 3D culture of G7m and E2 cells. RNAseq analysis was performed as previously described [9].

## Results

### Optimisation of the ClonoScreen3D platform

Five patient-derived GBM cell lines (MGMT promoter methylated G7s, G7m and E2, and MGMT promoter unmethylated S2 and GBML20) were used to evaluate the feasibility of converting the 3D CSA from 12-well format to 96-well format. G7s, G7m, and GBML20 cell lines formed distinct colonies with plating efficiencies of 30-50%, compared to diffuse growth and low plating efficiency (<20%) observed for E2 and S2 (Fig. 1A). Accordingly, further assay development was performed using G7s cells. Seeding density and colony growth times were optimized to obtain sufficient countable colonies under control and irradiated conditions. To validate the screen with radiation, G7s cells were exposed to 3 Gy (Fig. 1B), a dose selected based on survival responses to radiation alone and in combination with various compounds (rucaparib example shown in Fig. 1C). Colony forming ability and radiation/drug responses were not significantly affected by the transition to 96-well format.

**Figure 1.**
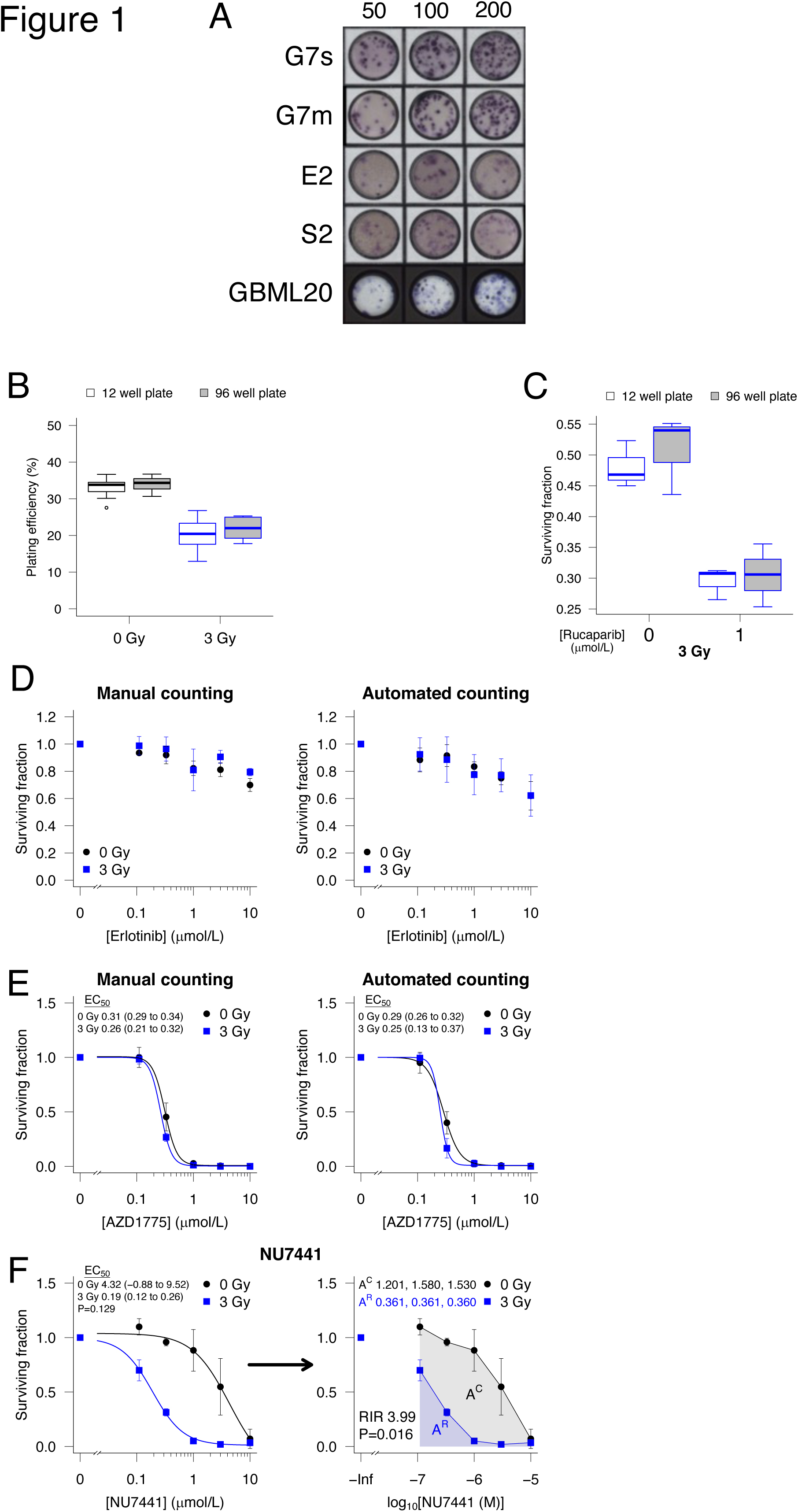
Optimisation of the ClonoScreen3D screening clonogenic assay format. **A**, Representative images of MTT-stained and PFA fixed colonies of patient-derived cell lines G7s, G7m, E2, S2, and GBML20 seeded in 96-well 3D-Alvetex plates under stem-enriched conditions at 50, 100 and 200 cells per well, incubated for 14 days. **B**, Clonogenic plating efficiency of G7s cells following sham irradiation or exposure to 3 Gy in 12-well and 96-well clonogenic assay format, *n*≥6. **C**, Clonogenic survival of G7s cells treated with 3 Gy alone or in combination with rucaparib (1 μmol/L) in 12-well and 96-well clonogenic assay format, *n*=6. Boxplots presented according to the Tukey method. **D**, Clonogenic survival of G7s cells following treatment with erlotinib or vehicle two hours prior to IR (3 Gy) calculated using manual or automated colony counting. **E**, Clonogenic survival of G7s cells following treatment with AZD1775 or vehicle two hours prior to IR (3 Gy) calculated using manual or automated colony counting. **F**, Clonogenic survival of G7s cells following treatment with NU7441 or vehicle two hours prior to IR (3 Gy) calculated from automated colony counts. The radiation interaction ratio (RIR) is calculated by comparison of areas-under-the-curve of the control (0 Gy) and irradiatied (3 Gy) samples, following log-transformation of concentration. The surviving fraction of irradiated samples was computed using the plating efficiency of the vehicle + 3 Gy control, normalizing for the effect of radiation alone. Points represent mean ± standard deviation (sd), *n*=3. EC_50_ (μmol/L) with 95% confidence interval calculated by fitting of a 4-parameter dose response curve. EC_50_ values compared by testing means of ratios.

To maximise efficiency, an automated colony counting process was developed using the opensource software packages ImageJ (https://imagej.net/software/fiji/) and OpenCFU [25] and compared against manual counting. Manual and automated counts showed a strong positive correlation, although some small differences in absolute values were noted (Spearman’s rho 0.91 P<0.001, Supp. Fig. S1). Importantly, using drug response as the critical endpoint, near identical responses to erlotinib (Fig. 1D) and AZD1775 (Fig. 1E) were observed, with or without radiation. These plots show data normalized for ionizing radiation (IR) effects, to highlight interactions between drug and radiation as shifts in drug response. Since only a single radiation dose was tested, we describe these plots as indicating ‘radiosensitizing potential’ rather than ‘radiosensitization’ *per se*, which generally requires multiple radiation dose points. Notably, the 96-well 3D-CSA enabled quantification of single agent activity (SAA) as well as IR interactions, thereby also informing selection of drug concentration(s) for further radiosensitization studies.

### Radiation Interaction Ratio (RIR) to quantify interaction between drugs and radiation

Initial characterization of ClonoScreen3D utilized NU7441, a known radiosensitizer that inhibits the key double strand break repair protein DNA-dependent protein kinase (DNA-PK) [26]. Whilst NU7441 showed weak SAA in the G7s cell line (Fig. 1F), it exhibited the expected marked interaction with IR, as indicated by the shift in drug dose response curve.

While SAA can be readily parameterized in terms of drug EC_50_, no analysis method has been developed to quantify the interaction with radiation. For drugs exhibiting SAA, comparison of EC_50_ values for drug alone and in combination with IR is indicative of radiosensitizing activity [27]. However, this approach is not applicable when drugs lack SAA or do not conform to a classical dose response, as in Fig. 1F (0 Gy), where EC_50_ cannot be estimated with meaningful confidence. In such situations, statistical comparison of EC_50_ values fails to capture even marked shifts in dose response (Fig. 1F P=0.129). To address this issue, we determined areaunder-the-curve (AUC) values for control and irradiated samples, an approach inspired by the widely used ‘mean inactivation dose’ parameter [28]. This generated a novel value, which we termed the ‘radiation interaction ratio’ (RIR), that is defined as the ratio of AUCs and captures the relative shift in dose response curves, thus providing a quantitative readout of radiosensitizing potential. In the example shown in Fig. 1F, the RIR was found to be 3.99 (P=0.016), confirming statistical significance and quantifying the observed interaction between NU7441 and IR.

### Quantification of radiosensitizing activity of PARP inhibitors using the ClonoScreen3D platform

Since the DNA repair enzyme poly(ADP-ribose) polymerase 1 (PARP-1) is overexpressed in GBM and shows very low expression in healthy brain tissue, it is a promising therapeutic target [29]. PARP inhibitors have consistently shown radiosensitizing effects in preclinical models of GBM both *in vitro* and *in vivo* [30–32] and are currently under investigation in phase I and II clinical trials [33].

To validate ClonoScreen3D as a radiosensitizer screening tool, we evaluated the radiation interactions of three PARP inhibitors (PARPi): rucaparib, talazoparib, and olaparib, the latter two known to radiosensitize GBM [13, 34]. Using ClonoScreen3D, these compounds all exhibited radiation interactions in G7s cells (Fig. 2A), with talazoparib having the highest RIR value (RIR 2.53 P<0.001), followed by rucaparib (RIR 1.95 P=0.002), and olaparib (RIR 1.56 P=0.003). Analysis by ClonoScreen3D also revealed that talazoparib exhibited potent SAA (EC_50_ 32 nmol/L, 95% CI 27 to 37 nmol/L) unlike the other PARP inhibitors tested to date.

**Figure 2.**
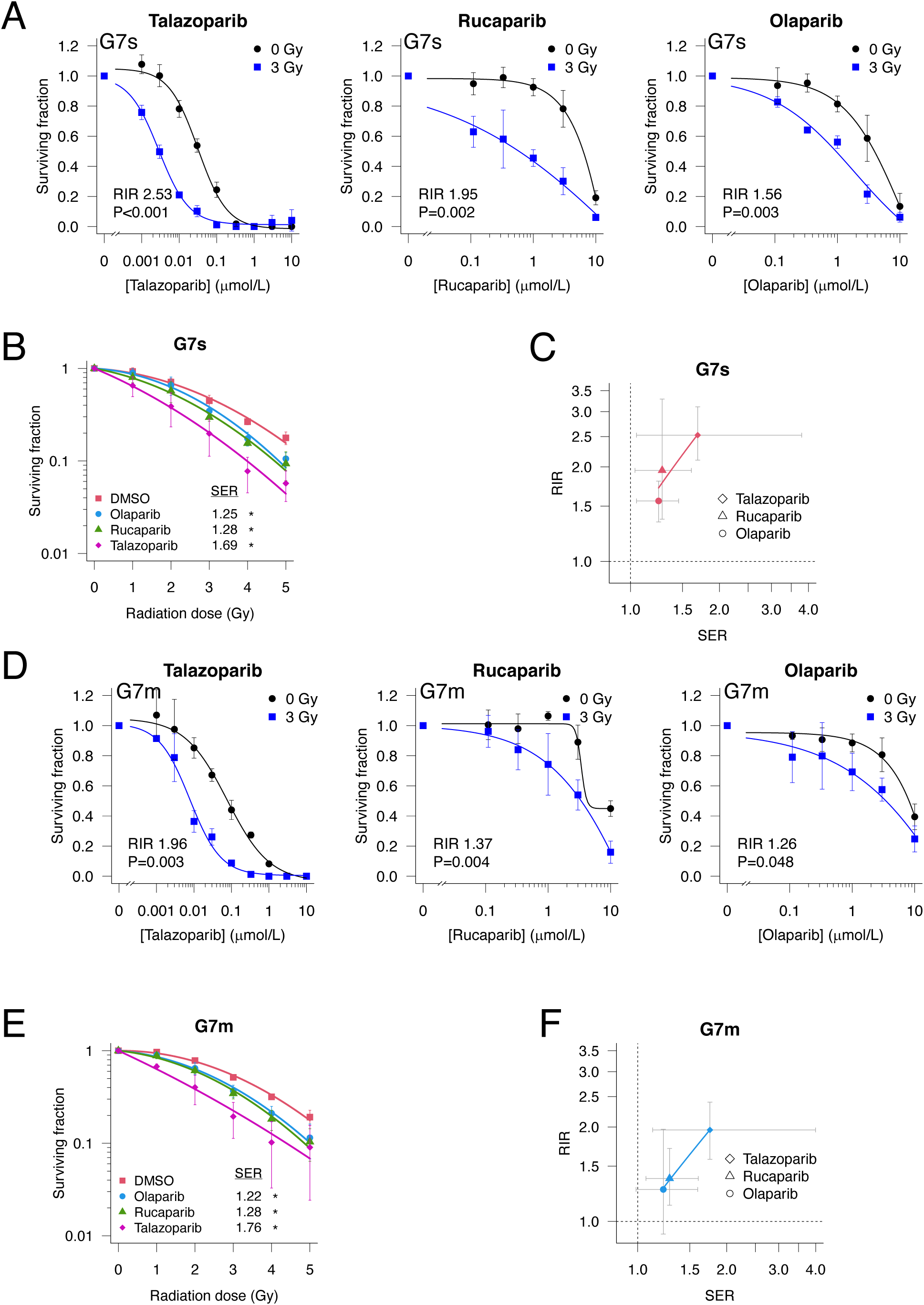
Validation of the ClonoScreen3D platform for identification of radiosensitizers using PARP inhibitors. **A**, Clonogenic survival of G7s cells following treatment with PARP inhibitors or vehicle two hours prior to IR (3 Gy). The surviving fraction of irradiated samples was computed using the plating efficiency of the vehicle + 3 Gy control, normalizing for the effect of radiation alone. Data fitted with a 4-parameter dose response curve. **B**, Full 3D radiation dose response clonogenic survival of G7s cells treated with olaparib (1 μmol/L), rucaparib (1 μmol/L) and talazoparib (5 nmol/L) two hours prior to IR. Data fitted using the linear quadratic model. Sensitizer enhancement ratios (SER) calculated using linear quadratic mean inactivation dose and subject to one-tailed ratio t-test. **C**, Correlation of RIR and SER values determined for PARP inhibitors in G7s cells. Points represent values calculated from three independent experiments, and error bars indicate coefficient 95% confidence intervals. **D**, Clonogenic survival of G7m cells following treatment with PARP inhibitors or vehicle two hours prior to IR (3 Gy). The surviving fraction of irradiated samples was computed using the plating efficiency of the vehicle + 3 Gy control, normalizing for the effect of radiation alone. Data fitted with a 4-parameter dose response curve. **E**, Full 3D radiation dose response clonogenic survival of G7m cells treated with olaparib (1 μmol/L), rucaparib (1 μmol/L) and talazoparib (5 nmol/L) two hours prior to IR. Data fitted using the linear quadratic model. Sensitizer enhancement ratios (SER) calculated using linear quadratic mean inactivation dose and subject to one-tailed ratio t-test. **F**, Correlation of RIR and SER values determined for PARP inhibitors in G7m cells. Points represent values calculated from three independent experiments, and error bars indicate coefficient 95% confidence intervals. Unless otherwise stated, points represent mean ± sd, *n*=3.

To confirm true radiosensitizing activity, PARPi were tested in 12-well 3D-CSA with multiple radiation dose (0 to 5 Gy). Based on ClonoScreen3D data, olaparib and rucaparib were dosed at 1 µmol/L and talazoparib at 5 nmol/L owing to its potent SAA. As expected, all three PARPi caused significant radiosensitization (sensitizer enhancement ratio, SER >1), confirming the ability of RIR to detect radiosensitizers (Fig. 2B, Supp. Table 2). Furthermore, RIR values exhibited a monotonic relationship with gold-standard SER in G7s cells, across the three PARPi tested (Fig. 2C).

To assess the generalizability of RIR to identify GBM radiosensitizers, we determined RIR values for PARPi in a second GBM model (G7m), which had been optimized for ClonoScreen3D (Supp. Fig. S2). Consistent with the previous result, talazoparib was the only PARPi to exhibit potent SAA (EC_50_ 76 nmol/L, 95% CI 42 to 110 nmol/L); and also showed the greatest interaction with IR (RIR 1.96 P=0.003), followed by rucaparib (RIR 1.37 P=0.004) and olaparib (RIR 1.26 P=0.048, Fig. 2D). Significant radiosensitization was observed in full CSA for each PARPi in G7m cells (Fig. 2E), which again correlated monotonically with RIR (Fig. 2F). Together, these findings demonstrate the utility of RIR for identification and prioritization of candidate radiosensitizers.

### Quantitative comparison of radiosensitizing effects of different DDR inhibitors

Although the radiosensitizing activities of multiple DDR inhibitors have been studied individually, direct comparison of inhibitors targeting different pathways has not been widely reported. We screened fifteen inhibitors targeting different DDR proteins using G7s cells and the ClonoScreen3D assay workflow summarized in Fig. 3. Five compounds exhibited potent SAA (EC_50_ < 2 µmol/L, Fig. 4A and B; Supp. Table 3), with talazoparib displaying the highest activity, followed by inhibitors targeting Chk1/2 and ATR. Compounds were ranked by RIR adjusted P-value (Fig. 4A), to encapsulate both the magnitude and reproducibility of the interaction with IR. In addition to summary graphics, individual drug response curves are presented in Supp. Fig. S3 and S4, and numerical results reported in Supp. Table 3 and 4.

**Figure 3.**
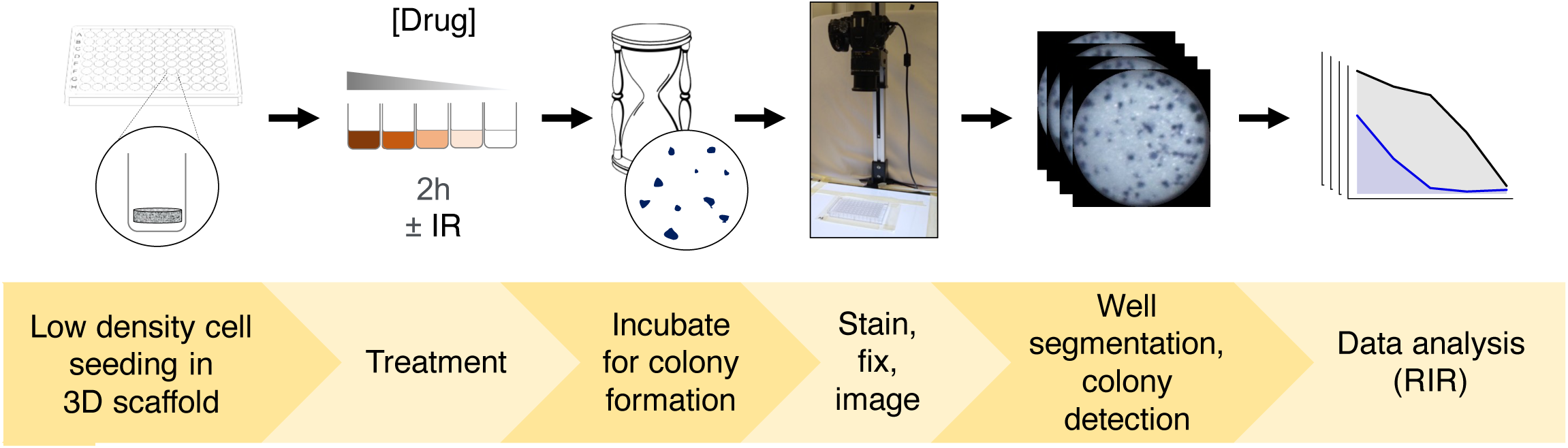
Schematic depicting the workflow for ClonoScreen3D when screening compounds for interaction with radiation. Graphical description of the methodology for the ClonoScreen3D platform. Cells are seeded, incubated for 18 hours and treated with respective compounds over the desired concentration range. After 2 hours incubation, cells are irradiated or sham-irradiated and incubated for colony formation. Colonies are stained with MTT, images of plates acquired with a digital camera followed by well segmentation using ImageJ and automated colony counting with OpenCFU. Following normalization for the effect of IR, radiation interaction ratios (RIR) are determined. Compound single agent activity is additionally quantified in terms of EC_50_, following fitting of a 4-parameter dose response model.

**Figure 4.**
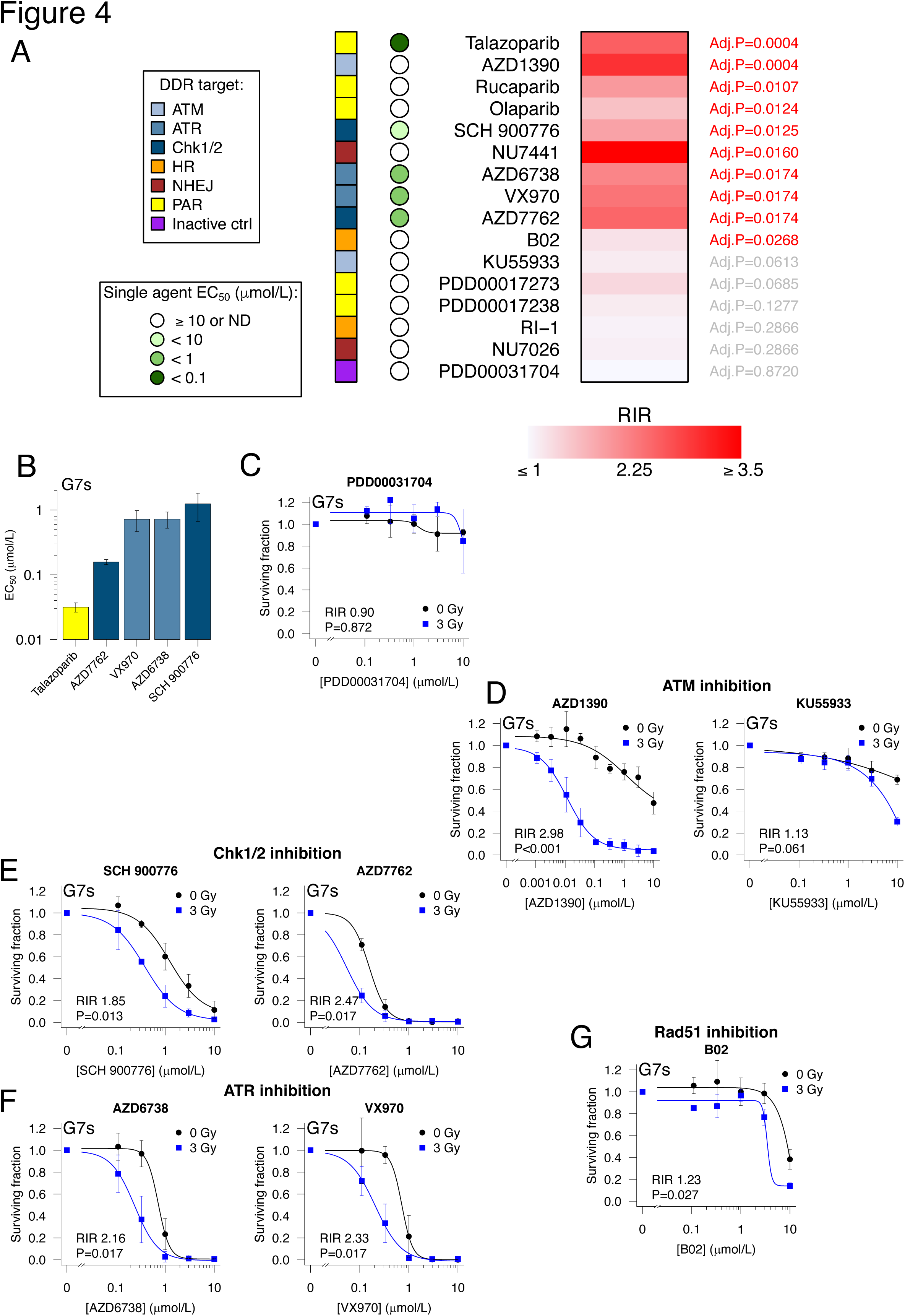
Quantitative comparison of the radiation interaction of DNA damage response inhibitors using the ClonoScreen3D platform. **A**, A panel of DNA damage response (DDR) inhibitors was screened for interaction with radiation using the ClonoScreen3D platform in G7s cells. Cells were incubated with drugs for two hours prior to irradiation (3 Gy). Radiation interaction ratio (RIR) values were computed and compounds ranked by FDR adjusted P-value, following one-tailed ratio t-testing. Drug single agent activity was quantified as EC_50_ following fitting of a 4-parameter dose response model to sham-irradiated samples. The target pathway or protein is indicated. HR homologous recombination, NHEJ non-homologous end-joining, PAR poly(ADP-ribose), ND not determined. Data generated in three independent experiments. **B**, DDR inhibitors demonstrating single agent activity in G7s cells. Bars represent EC_50_ with error bars indicating 95% confidence interval. **C**, Clonogenic survival of G7s cells treated with an inactive control compound, PDD00031704, and IR (3 Gy). **D**, Clonogenic survival of G7s cells treated with ATM inhibitors, and IR (3 Gy). **E**, Clonogenic survival of G7s cells treated with Chk1/2 inhibitors, and IR (3 Gy). **F**, Clonogenic survival of G7s cells treated with ATR inhibitors, and IR (3 Gy). **G**, Clonogenic survival of G7s cells treated with the homologous recombination inhibitor B02, and IR (3 Gy). The surviving fraction of irradiated samples was computed using the plating efficiency of the vehicle + 3 Gy control, normalizing for the effect of radiation alone. Data fitted with a 4-parameter dose response curve. Points represent mean ± sd, *n*=3.

To assess the specificity of the assay, an inactive control compound was included. PDD00031704 is a modified variant of a PARG inhibitor, with on-target IC_50_ >100 µmol/L. In the ClonoScreen3D assay, this compound ranked lowest for interaction with radiation in G7s and G7m cells (Fig. 4A and C; Supp. Fig. S4). The ATM inhibitor AZD1390 exhibited a marked interaction with IR (RIR 2.98 P<0.001), while the less potent ATM inhibitor KU55933 showed limited (non-significant) activity only at 10 µmol/L (Fig. 4D). The next highest ranked compound was SCH 900776, an inhibitor of Chk1/2 (RIR 1.85 P=0.013, Fig. 4E), while another Chk1/2 inhibitor AZD7762 was also identified as a hit (RIR 2.47 P=0.017). Two ATR inhibitors, AZD6738 (RIR 2.16 P=0.017) and VX970 (RIR 2.33 P=0.017), were also identified as interacting significantly with IR (Fig. 4F). The DNA-PK inhibitor NU7441 exhibited the greatest absolute interaction with IR (RIR 3.99 P=0.016, Fig. 1F), although its ranking was reduced by inter-replicate variability. By contrast, a second DNA-PK inhibitor, NU7026 showed no significant activity. Two homologous recombination (HR)-targeting inhibitors exhibited limited radiation interaction, which was statistically significant only for B02 (RIR 1.23 P=0.017, Fig. 4G); this may be explained by their low drug potencies, having EC_50_ values in the micromolar range.

Of the fifteen DDR inhibitors tested, eight compounds showed a significant interaction with radiation in both G7s and G7m cells (Fig. 4A and D-G, Supp. Fig. SF4). A further five compounds showed radiosensitizing potential in a single cell line.

### Identification of novel targets for radiosensitization of GBM cells

To identify novel radiosensitization targets, we performed RNAseq analysis on samples obtained from two patient-derived GBM cell lines, E2 and G7m, that had been cultured in 3D and treated with IR (5 Gy) or sham-irradiated four hours previously. The timepoint of RNA extraction was chosen to identify early response genes regulating radioresistance in GBM. Since only modest changes in gene expression were observed in E2 cells, we focused on G7m cells. In this context, exposure to IR significantly altered expression of multiple DDR genes (e.g. BRCA1, RAD51, XRCC2) as expected, and also included genes associated with: (*i*) reduced cell cycle (CDC25C, CDKN1B, CDK18) and mitotic progression (BUB1, PLK1, CDC25A); (*ii*) NFκB signalling (NFKBIE, NFKBIA and NFKB2); (*iii*) cytokine and growth factor signalling (CXCL10, CXCL6, VEGFA); and (*iv*) stem cell regulation (FOS, WNT2, WNT5A, ID4) (Fig. 5A). A full list is provided in Supplementary File 1.

**Figure 5.**
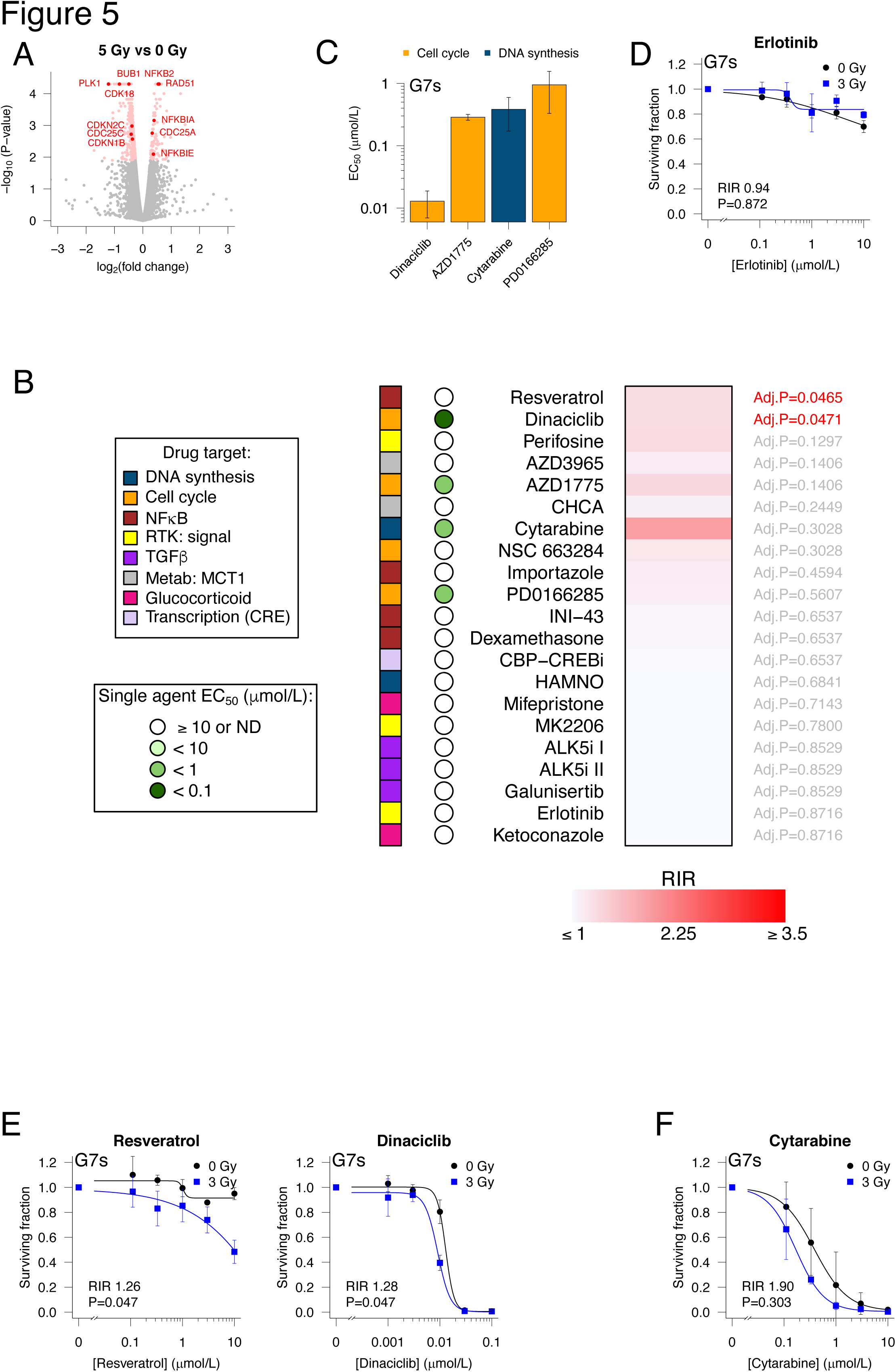
Evaluation of drug-radiation combinations using the ClonoScreen3D platform for identification of novel radiosensitizers. **A**, Transcriptomic changes in G7m cells cultured in 3D conditions were measured four hours after exposure to 5 Gy. Selected significantly upregulated and downregulated genes of interest are annotated, *n*=3. **B**, A panel of commercially available inhibitors targeting pathways identified by transcriptomic analysis was screened for interaction with radiation using the ClonoScreen3D platform in G7s cells. Cells were incubated with drugs for two hours prior to irradiation (3 Gy). Radiation interaction ratio (RIR) values were computed and compounds ranked by FDR adjusted P-value, following one-tailed ratio t-testing. Drug single agent activity was quantified as EC_50_ following fitting of a 4-parameter dose response model to sham-irradiated samples. The target pathway or protein is indicated. DDR DNA damage response, RTK receptor tyrosine kinase, Metab metabolism, CRE cAMP response element. Data generated in three independent experiments. **C**, Inhibitors demonstrating single agent activity in G7s cells. Bars represent EC_50_ with error bars indicating 95% confidence interval. **D**, Clonogenic survival of G7s cells treated with erlotinib and IR (3 Gy). **E**, Clonogenic survival of G7s cells treated with dinaciclib or resveratrol, and IR (3 Gy). **F**, Clonogenic survival of G7s cells treated with cytarabine and IR (3 Gy). The surviving fraction of irradiated samples was computed using the plating efficiency of the vehicle + 3 Gy control, normalizing for the effect of radiation alone. Data fitted with a 4-parameter dose response curve. Points represent mean ± sd, *n*=3.

Commercially available inhibitors targeting pathways and gene products identified in this experiment were selected for evaluation as potential radiosensitizers using ClonoScreen3D. Where several genes from the same biological pathway were identified, additional compounds targeting the pathway were included, such as the WEE1 inhibitor AZD1775 to target mitosis and the cyclin-dependent kinase (CDK) inhibitor dinaciclib for cell cycle progression.

In this prospective screen, inhibitors targeting cell cycle regulation, in particular the G2/M checkpoint, demonstrated marked SAA in G7s cells (Fig. 5B and C, Supp. Table 5, Supp. Fig. S5). Dinaciclib, a potent small molecule inhibitor of CDK1, CDK2, CKD5 and CDK9, exhibited the highest SAA (EC_50_ 13 nmol/L, 95% CI 4 to 21 nmol/L). Inhibitors targeting mitotic progression (AZD1775, PD0166285) also showed potent SAA as did cytarabine, a pyrimidine nucleoside analogue that inhibits DNA synthesis [9, 35]. A similar pattern of SAA was observed in G7m cells (Supp. Fig. S6, Supp. Table 6).

### Targeting DNA synthesis and cyclin-dependent kinases induces radiosensitization of GBM cells

In keeping with previous 3D CSA and clinical trials [9, 35] erlotinib exhibited no SAA or radiosensitizing activity in G7s (RIR 0.94 P=0.872; Fig. 5D) or G7m cells (RIR 1.06 P=0.104; Supp. Fig. S6). Of the compounds targeting radiation-response processes identified by transcriptomic analysis, only two exhibited a robust interaction with radiation, following correction of P-values for multiple comparison. Resveratrol, an NFκB inhibitor, demonstrated a modest interaction with IR in G7s cells (RIR 1.26 P=0.047, Fig. 5E). In addition to potent SAA, dinaciclib exhibited an interaction with IR in G7s cells (RIR 1.28 P=0.047). Despite failing to achieve statistical significance due to inter-replicate variability, the DNA synthesis inhibitor cytarabine exhibited the highest RIR value in this screen, in both G7s and G7m cells (G7s RIR 1.90 P=0.303; G7m RIR 1.76 P=0.104, Fig. 5F, Supp. Fig. S6). Because the RIR magnitude was high, the interaction with IR was additionally tested by statistical comparison of EC_50_ values between the combination and radiation-only samples. A significant reduction in EC_50_ was observed when cytarabine was combined with IR in G7m (P=0.017) but not G7s (P=0.119) cells (Supp. Fig. S7). This result and the SAA of cytarabine justified its selection for further investigation.

To confirm the ability of the ClonoScreen3D platform to identify *bona fide* novel radiosensitizers, gold-standard full radiation dose response CSAs (12-well) were performed for the novel hit compounds dinaciclib and cytarabine. Dinaciclib significantly radiosensitized G7s (SER 1.47 P=0.010), G7m (SER 1.25 P=0.002) and E2 (SER 1.55 P=0.001) cells at 10 nmol/L, with E2 cells also radiosensitized at 1 nmol/L (SER 1.17 P=0.031, Fig. 6A and Suppl. Table 7).

**Figure 6.**
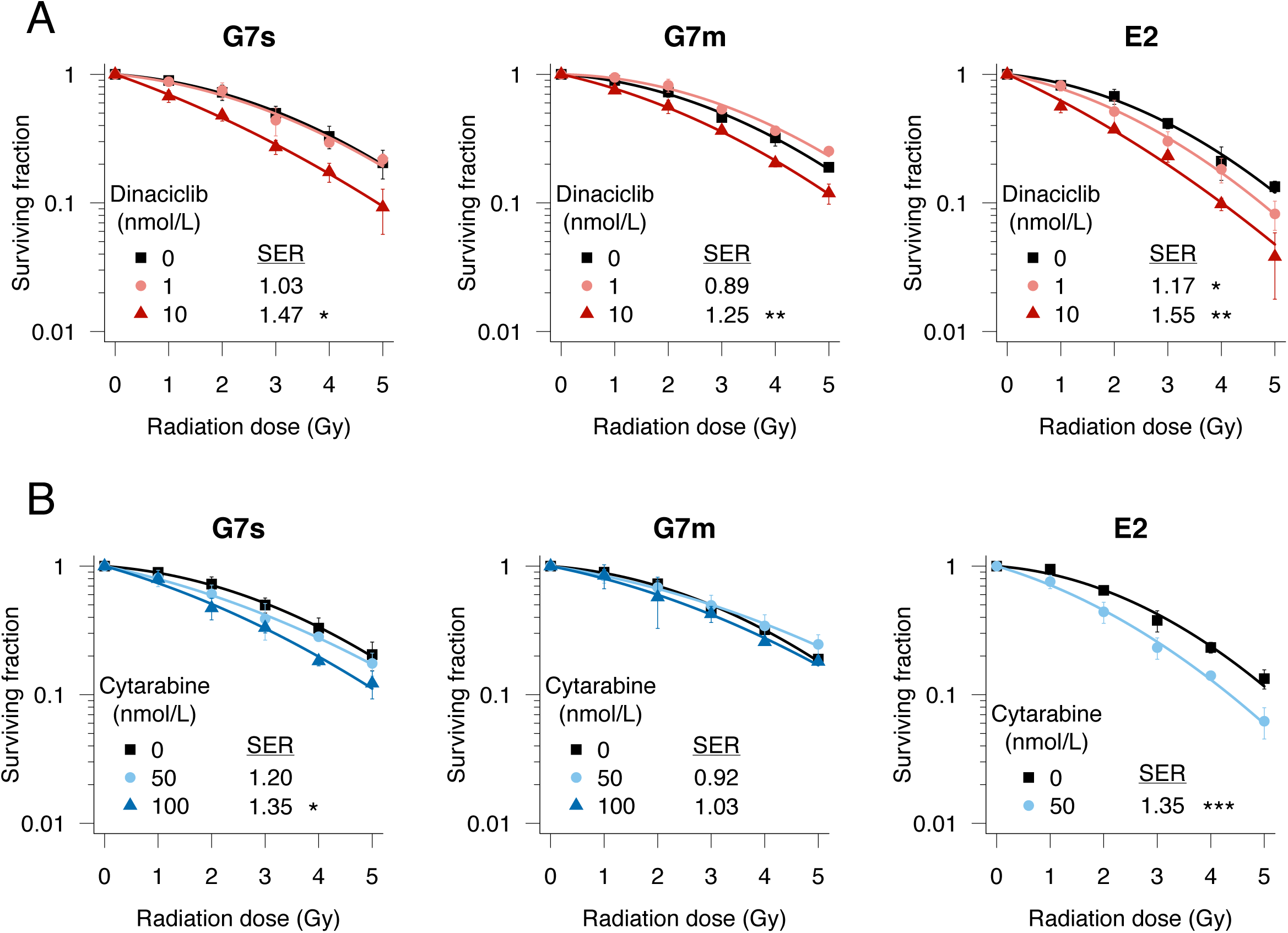
Dinaciclib and cytarabine exhibit radiosensitizing activity in GBM cells. **A**, Full 3D radiation dose response clonogenic survival of G7s, G7m and E2 GBM cells treated with dinaciclib (1 and 10 nmol/L) two hours prior to IR. **B**, Full 3D radiation dose response clonogenic survival of G7s, G7m and E2 GBM cells treated with cytarabine (50 and 100 nmol/L) two hours prior to IR. Data fitted using the linear quadratic model. Sensitizer enhancement ratios (SER) calculated using linear quadratic mean inactivation dose and subject to one-tailed ratio t-test. Points represent mean ± sd, *n*=3.

Significant radiosensitization was also elicited by cytarabine (100 nmol/L) in G7s (SER 1.35 P=0.020) but not G7m cells (Fig. 6B). E2 cells were significantly radiosensitized by cytarabine at 50 nmol/L (SER 1.35 P=0.001). These studies confirmed concentration-dependent radiosensitizing activity of dinaciclib and cytarabine in primary GBM cells cultured in 3D.

## Discussion

Preclinical research is urgently required to identify therapeutic strategies for GBM that will translate into the clinic. Radiotherapy is a mainstay GBM treatment but does not achieve cure, and radiation dose-escalation is prohibited by lack of efficacy and normal brain toxicity [36]. The development of combination strategies to selectively sensitize GBM cells to IR is an attractive approach but has been severely limited by the low-throughput and/or low-fidelity of the preclinical models currently available.

We have successfully refined an *in vitro* 3D model of GBM with demonstrable clinical-relevance for use as a screening platform for novel radiotherapy-drug combinations, with goldstandard clonogenic survival as the readout. This is a major advance on existing screens involving IR, which generally rely on cell proliferation or short-term viability as readouts [37]. These assays have limitations: (*i*) viability readouts fail to discriminate cells that have lost their capacity to reproduce indefinitely; (*ii*) they are performed at early time-points, measuring responses after only one or two mitoses, whereas radiation-induced damaged cells may divide several times before becoming succumbing to reproductive death. The CSA is the only *bona fide* long-term reproductive integrity assay, and hence the most clinically-relevant.

Increasing the throughput of radiosensitizer identification experiments required reformatting of the CSA. Importantly, conversion of the ClonoScreen3D platform to 96-well format did not significantly influence treatment responses – a limitation reported previously [15, 38]. This enabled the platform to compare a broad range of DDR inhibitors as well as enabling screening of expansive drug libraries in 3D cultures of primary GBM cells, for this first time.

A novel analytical parameter, the RIR, was developed based on established radiobiological approaches to enable flexible, reproducible and statistically quantifiable assessment of the interactions between radiation and drugs, even when drug activity is not known *a priori*. Although modestly overestimating magnitude (reducing the likelihood of false-negative results), the RIR values for PARPi correlated with SER values determined using full CSA, confirming the utility of this novel parameter. In contrast to EC_50_ comparison, well-defined SAA drug activity is not required for RIR computation, rendering it highly advantageous for screening applications. This parameter accurately predicted activity of known radiosensitizers [39] and discriminated non-active compounds (e.g. PDD00031704 [40], erlotinib [9, 35]).

Pragmatically, the relative probabilities of false-negative and false-positive findings can be tuned through P-value adjustment methodology. For example, the significance of dinaciclib’s interaction with radiation was not maintained after P-value correction in G7m cells, but it was subsequently validated as a true radiosensitizer. To ensure robustness, hits identified in the ClonoScreen3D assay should be subject to full radiation dose CSA, to confirm and quantify radiosensitizing activity.

A recently published 2D CSA screening methodology included an established GBM cell [15]. Comparison of this system with ClonoScreen3D suggested several advantages of our platform, in addition to the notable improvement in clinical relevance offered by 3D culture [9, 41]. The method of Gomes *et al.* required viral transduction for colony detection, which may have consequences for cell behaviour. The assay also used high radiation doses (9 Gy) and a single drug concentration, meaning that narrow therapeutic combination windows may be missed. Furthermore, a secondary round of screening was required to distinguish between single agent and combination activity, whereas ClonoScreen3D quantifies both simultaneously and informs on drug concentrations for validation experiments. Lastly, the use of RIR to quantify drug-radiation interactions may be more intuitive and familiar to radiation biologists or oncologists than Z-score based metrics.

Given their critical role in the radiation response, inhibition of DDR proteins is expected to potentiate treatment efficacy [42]. As quantified by RIR, 13/15 (87%) of the DDR inhibitors tested showed a significant interaction with IR in at least one GBM cell context, confirming the assay’s ability to detect potential radiosensitizing activity. The ClonoScreen3D platform also confirmed therapeutic interactions mediated by drugs targeting other pathways previously suggested to radiosensitize GBM cells, albeit in 2D conditions, for example NFκB signalling (resveratrol) [43].

ClonoScreen3D also identified novel radiosensitizing compounds including the FDA approved compound dinaciclib. Dinaciclib is a selective and potent inhibitor of CDK1, CDK2, CDK5, and CDK9 with EC_50_ values of 1-4 nmol/L across the target CDKs [44]. This compound has been well tolerated in clinical trials, exhibiting efficacy in patients with chronic lymphocytic leukemia [45] and relapsed multiple myeloma [46]. Consistent with our findings, SAA of dinaciclib has been previously reported in 2D and 3D GBM cells [47, 48]. A second FDA-approved drug, cytarabine, was also identified as a radiosensitizer. Cytarabine has exhibited clinical activity in two patients with astrocytoma following intraventricular administration of a liposomal formulation [49]. Regrettably, this study was terminated prematurely because of slow recruitment. Crucially, following identification by ClonoScreen3D, dinaciclib and cytarabine were validated as true radiosensitizers in the gold-standard radiation dose response 3D CSA.

The work presented here describes a novel drug-screening methodology for identification of radiosensitizers using cell culture technology that reproduces clinical treatment responses. This technology allowed comparison of multiple compounds undergoing evaluation in clinical trials. Our findings support the notion that targeting the DDR is likely to provide multiple opportunities for radiosensitization of GBM, for example with inhibitors of PARP (talazoparib), ATM (AZD1390), DNA-PK (NU7441) and Chk1 (SCH900776 and AZD7762), as long as they exhibit sufficient tumour penetration and do not exacerbate neurotoxicity.

Inhibitors of other DDR proteins, for example PARG, interacted with IR in a cell contextdependent manner, confirming the need to validate compounds in multiple primary cell lines. Finally, the paucity of translatable inhibitors of HR repair [50] was highlighted by the screen. The majority of non-DDR targeting drugs included in our initial screen showed little radiosensitizing potential highlighting the strategic importance of DDR modulators. Advantageously, ClonoScreen3D allows newly identified candidate drugs to be benchmarked against established radiosensitizers.

The current dismal prognosis despite aggressive treatment suggests that novel combination approaches will have a crucial role in GBM therapy. Our results validate the ClonoScreen3D assay platform for identification and comparison of novel radiosensitizers for GBM and has clear potential for extension to other cancer types. This assay provides a new technology that will underpin an improved drug development pipeline, accelerating development of lead compounds to augment the efficacy of radiotherapy.

## Acknowledgements

This work was funded by a NC3Rs grant (Grant Ref: NC/P001335/1) awarded to N. Gomez-Roman and A.J.Chalmers. Additional support was provided by the Cancer Research UK Radiation Research Centre of Excellence at the University of Glasgow (C16583/A28803). We would like to thank Prof. Colin Watts (University of Birmingham) and Dr Dimitris Placantonakis (NYU) who kindly donated cell lines.

## References

1. Louis, D.N., et al., The 2016 World Health Organization Classification of Tumors of the Central Nervous System: a summary. Acta Neuropathol, 2016. 131(6): p. 803–20.

2. Stupp, R., et al., Effects of radiotherapy with concomitant and adjuvant temozolomide versus radiotherapy alone on survival in glioblastoma in a randomised phase III study: 5-year analysis of the EORTC-NCIC trial. Lancet Oncology, 2009. 10(5): p. 459–466.

3. Taylor, O.G., J.S. Brzozowski, and K.A. Skelding, Glioblastoma Multiforme: An Overview of Emerging Therapeutic Targets. Front Oncol, 2019. 9: p. 963.

4. Lassman, A.B., L.E. Abrey, and M.R. Gilbert, Response of glioblastomas to EGFR kinase inhibitors. The New England journal of medicine, 2006. 354(5): p. 525–6; author reply 525-6.

5. Lai, A., et al., Phase II study of bevacizumab plus temozolomide during and after radiation therapy for patients with newly diagnosed glioblastoma multiforme. Journal of clinical oncology: official journal of the American Society of Clinical Oncology, 2011. 29(2): p. 142–8.

6. Reardon, D.A., et al., Cilengitide: an RGD pentapeptide alphanubeta3 and alphanubeta5 integrin inhibitor in development for glioblastoma and other malignancies. Future Oncol, 2011. 7(3): p. 339–54.

7. Vogelbaum, M.A., et al., Response rate to single agent therapy with the EGFR tyrosine kinase inhibitor erlotinib in recurrent glioblastoma multiforme: Results of a phase II study. Neuro-Oncology, 2004. 6(4): p. 384–384.

8. Mason, W.P., End of the road: confounding results of the CORE trial terminate the arduous journey of cilengitide for glioblastoma. Neuro Oncol, 2015. 17(5): p. 634–5.

9. Gomez-Roman, N., et al., A novel 3D human glioblastoma cell culture system for modeling drug and radiation responses. Neuro Oncol, 2016.

10. Caragher, S., A.J. Chalmers, and N. Gomez-Roman, Glioblastoma’s Next Top Model: Novel Culture Systems for Brain Cancer Radiotherapy Research. Cancers (Basel), 2019. 11(1).

11. Walker, M.D., et al., Randomized Comparisons of Radiotherapy and Nitrosoureas for the Treatment of Malignant Glioma after Surgery. New England Journal of Medicine, 1980. 303(23): p. 1323–1329.

12. Bao, S., et al., Glioma stem cells promote radioresistance by preferential activation of the DNA damage response. Nature, 2006. 444(7120): p. 756–60.

13. Ahmed, S.U., et al., Selective Inhibition of Parallel DNA Damage Response Pathways Optimizes Radiosensitization of Glioblastoma Stem-like Cells. Cancer Res, 2015. 75(20): p. 4416–28.

14. Ye, R., et al., High-Content Clonogenic Survival Screen to Identify Chemoradiation Sensitizers. Int J Radiat Oncol Biol Phys, 2021. 111(5): p. e27–e37.

15. Gomes, N.P., et al., A High Throughput Screen with a Clonogenic Endpoint to Identify Radiation Modulators of Cancer. Radiat Res, 2023. 199(2): p. 132–147.

16. Tiwana, G.S., et al., Identification of vitamin B1 metabolism as a tumor-specific radiosensitizing pathway using a high-throughput colony formation screen. Oncotarget, 2015. 6(8): p. 5978–5989.

17. Katz, D., et al., Increased efficiency for performing colony formation assays in 96-well plates: novel applications to combination therapies and high-throughput screening. Biotechniques, 2008. 44(2): p. Ix–Xiv.

18. Lin, S.H., et al., A High Content Clonogenic Survival Drug Screen Identifies MEK Inhibitors as Potent Radiation Sensitizers for KRAS Mutant Non-Small-Cell Lung Cancer. Journal of Thoracic Oncology, 2014. 9(7): p. 965–973.

19. Al-Mayhani, T.M.F., et al., An efficient method for derivation and propagation of glioblastoma cell lines that conserves the molecular profile of their original tumours. Journal of neuroscience methods, 2009. 176(2): p. 192–199.

20. Ekstrøm, C.T., MESS: Miscellaneous Esoteric Statistical Scripts. 2019.

21. Schaarschmidt, G.D.a.M.H.a.D.G.a.F., mratios: Ratios of Coefficients in the General Linear Model. 2018.

22. Ritz, C., et al., Dose-Response Analysis Using R. PLoS One, 2015. 10(12): p. e0146021.

23. Ekstrøm, C.T., MESS: Miscellaneous Esoteric Statistical Scripts. 2019.

24. Gemechis Dilba Djira, M.H., Daniel Gerhard, Frank Schaarschmidt, mratios: Ratios of Coefficients in the General Linear Model. 2018.

25. Geissmann, Q., OpenCFU, a new free and open-source software to count cell colonies and other circular objects. PLoS One, 2013. 8(2): p. e54072.

26. Fang, X.G., et al., Inhibiting DNA-PK induces glioma stem cell differentiation and sensitizes glioblastoma to radiation in mice. Science Translational Medicine, 2021. 13(600).

27. Jackson, M.R., et al., Mesothelioma Cells Depend on the Antiapoptotic Protein Bcl-xL for Survival and Are Sensitized to Ionizing Radiation by BH3-Mimetics. Int J Radiat Oncol Biol Phys, 2020. 106(4): p. 867–877.

28. Subiel, A., R. Ashmore, and G. Schettino, Standards and Methodologies for Characterizing Radiobiological Impact of High-Z Nanoparticles. Theranostics, 2016. 6(10): p. 1651–1671.

29. Hanna, C., et al., Pharmacokinetics, safety and tolerability of olaparib and temozolomide for recurrent glioblastoma: results of the phase I OPARATIC trial. Neuro Oncol, 2020.

30. Chalmers, A.J., Overcoming resistance of glioblastoma to conventional cytotoxic therapies by the addition of PARP inhibitors. Anticancer Agents Med Chem, 2010. 10(7): p. 520–33.

31. Jannetti, S.A., et al., PARP-1-Targeted Radiotherapy in Mouse Models of Glioblastoma. J Nucl Med, 2018. 59(8): p. 1225–1233.

32. Venere, M., et al., Therapeutic targeting of constitutive PARP activation compromises stem cell phenotype and survival of glioblastoma-initiating cells. Cell Death Differ, 2014. 21(2): p. 258–69.

33. Sim, H.W., E. Galanis, and M. Khasraw, PARP Inhibitors in Glioma: A Review of Therapeutic Opportunities. Cancers (Basel), 2022. 14(4).

34. Lesueur, P., et al., Radiosensitization Effect of Talazoparib, a Parp Inhibitor, on Glioblastoma Stem Cells Exposed to Low and High Linear Energy Transfer Radiation. Sci Rep, 2018. 8(1): p. 3664.

35. Gomez-Roman, N., et al., Radiation Responses of 2D and 3D Glioblastoma Cells: A Novel, 3D-specific Radioprotective Role of VEGF/Akt Signaling through Functional Activation of NHEJ. Mol Cancer Ther, 2020. 19(2): p. 575–589.

36. Greene-Schloesser, D., et al., Radiation-induced brain injury: A review. Front Oncol, 2012. 2: p. 73.

37. Willers, H., et al., Screening and Validation of Molecular Targeted Radiosensitizers. Int J Radiat Oncol Biol Phys, 2021. 111(5): p. e63–e74.

38. Esquer, H., et al., Advanced High-Content-Screening Applications of Clonogenicity in Cancer. Slas Discovery, 2020. 25(7): p. 734–743.

39. Gorte, J., et al., Comparative Proton and Photon Irradiation Combined with Pharmacological Inhibitors in 3D Pancreatic Cancer Cultures. Cancers (Basel), 2020. 12(11).

40. James, D.I., et al., First-in-Class Chemical Probes against Poly(ADP-ribose) Glycohydrolase (PARG) Inhibit DNA Repair with Differential Pharmacology to Olaparib. ACS Chem Biol, 2016. 11(11): p. 3179–3190.

41. Storch, K., et al., Three-dimensional cell growth confers radioresistance by chromatin density modification. Cancer Res, 2010. 70(10): p. 3925–34.

42. Rominiyi, O. and S.J. Collis, DDRugging glioblastoma: understanding and targeting the DNA damage response to improve future therapies. Mol Oncol, 2022. 16(1): p. 11–41.

43. Wang, L., et al., Resveratrol, a potential radiation sensitizer for glioma stem cells both in vitro and in vivo. Journal of Pharmacological Sciences, 2015. 129(4): p. 216–225.

44. Parry, D., et al., Dinaciclib (SCH 727965), a novel and potent cyclin-dependent kinase inhibitor. Mol Cancer Ther, 2010. 9(8): p. 2344–53.

45. Flynn, J., et al., Dinaciclib is a novel cyclin-dependent kinase inhibitor with significant clinical activity in relapsed and refractory chronic lymphocytic leukemia. Leukemia, 2015. 29(7): p. 1524–9.

46. Kumar, S.K., et al., Dinaciclib, a novel CDK inhibitor, demonstrates encouraging single-agent activity in patients with relapsed multiple myeloma. Blood, 2015. 125(3): p. 443–8.

47. Jane, E.P., et al., Dinaciclib, a Cyclin-Dependent Kinase Inhibitor Promotes Proteasomal Degradation of Mcl-1 and Enhances ABT-737-Mediated Cell Death in Malignant Human Glioma Cell Lines. J Pharmacol Exp Ther, 2016. 356(2): p. 354–65.

48. Riess, C., et al., Cyclin-dependent kinase inhibitors exert distinct effects on patient-derived 2D and 3D glioblastoma cell culture models. Cell Death Discov, 2021. 7(1): p. 54.

49. Frankel, B.M., et al., Targeting Subventricular Zone Progenitor Cells with Intraventricular Liposomal Encapsulated Cytarabine in Patients with Secondary Glioblastoma: A Report of Two Cases. SN Compr Clin Med, 2020. 2(6): p. 836–843.

50. Durant, S.T., et al., The brain-penetrant clinical ATM inhibitor AZD1390 radiosensitizes and improves survival of preclinical brain tumor models. Sci Adv, 2018. 4(6): p. eaat1719.

