## Supplementary Figures for "ClonoScreen3D: a novel three-dimensional clonogenic screening platform for identification of radiosensitizers for glioblastoma"

### Supp. Figure S1

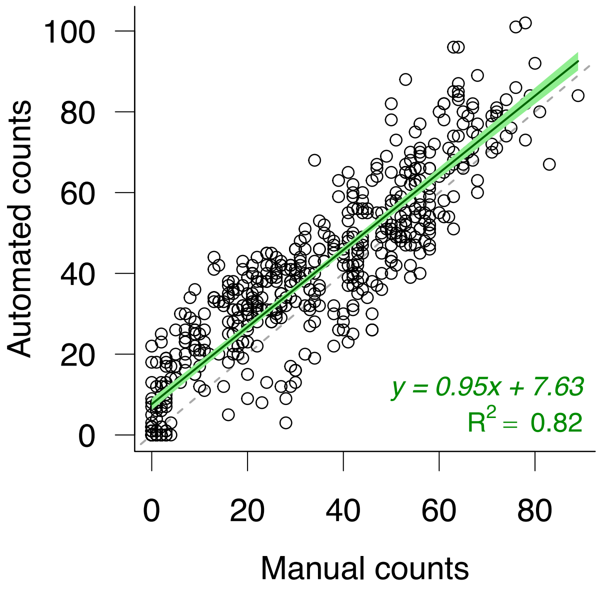


**Supp. Fig. S1: Correlation of manual and automated colony counts in G7s cells.** Colonies resulting from G7s cells treated with IR (3 Gy) or sham-IR in combination with vehicle, AZD1390 or erlotinib (0.0011 to 10 µmol/L) were counted manually and with an automated system (OpenCFU). Colony counts were determined from three independent experiments.

Supp. Figure S2

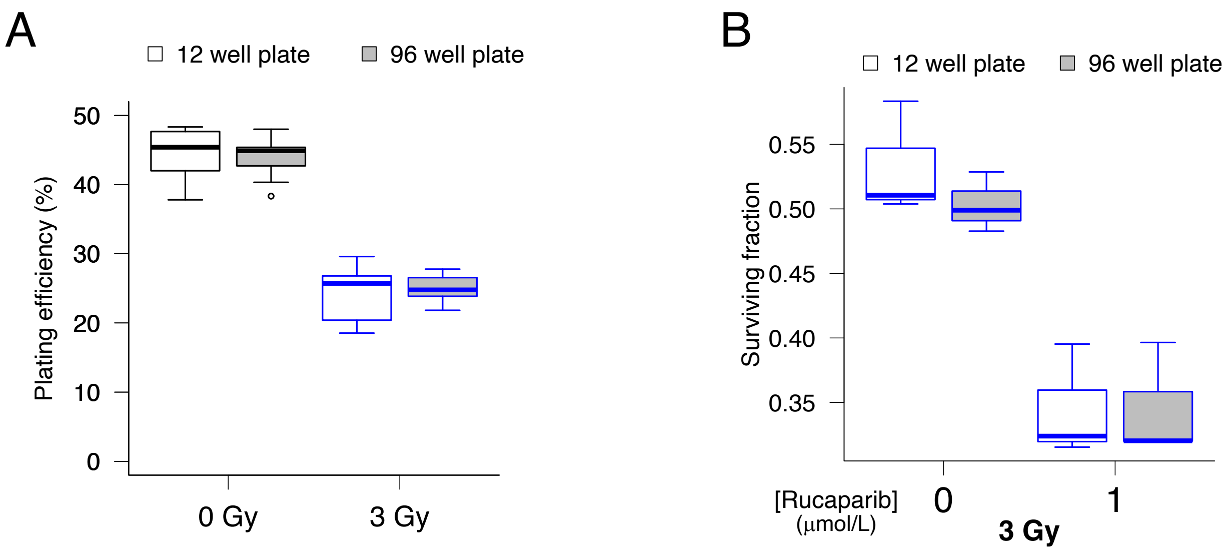


**Supp. Fig. S2: Clonogenic assay miniaturization in G7m cells.**

**A**, Clonogenic plating efficiency of G7m cells following sham irradiation or exposure to 3 Gy in 12-well and 96-well clonogenic assay format, *n*≥10. B, Clonogenic survival of G7m cells treated with 3 Gy alone or in combination with rucaparib (1 μmol/L) in 12-well and 96-well clonogenic assay format, *n*=3. Boxplots presented according to the Tukey method.

Supp. Figure S3

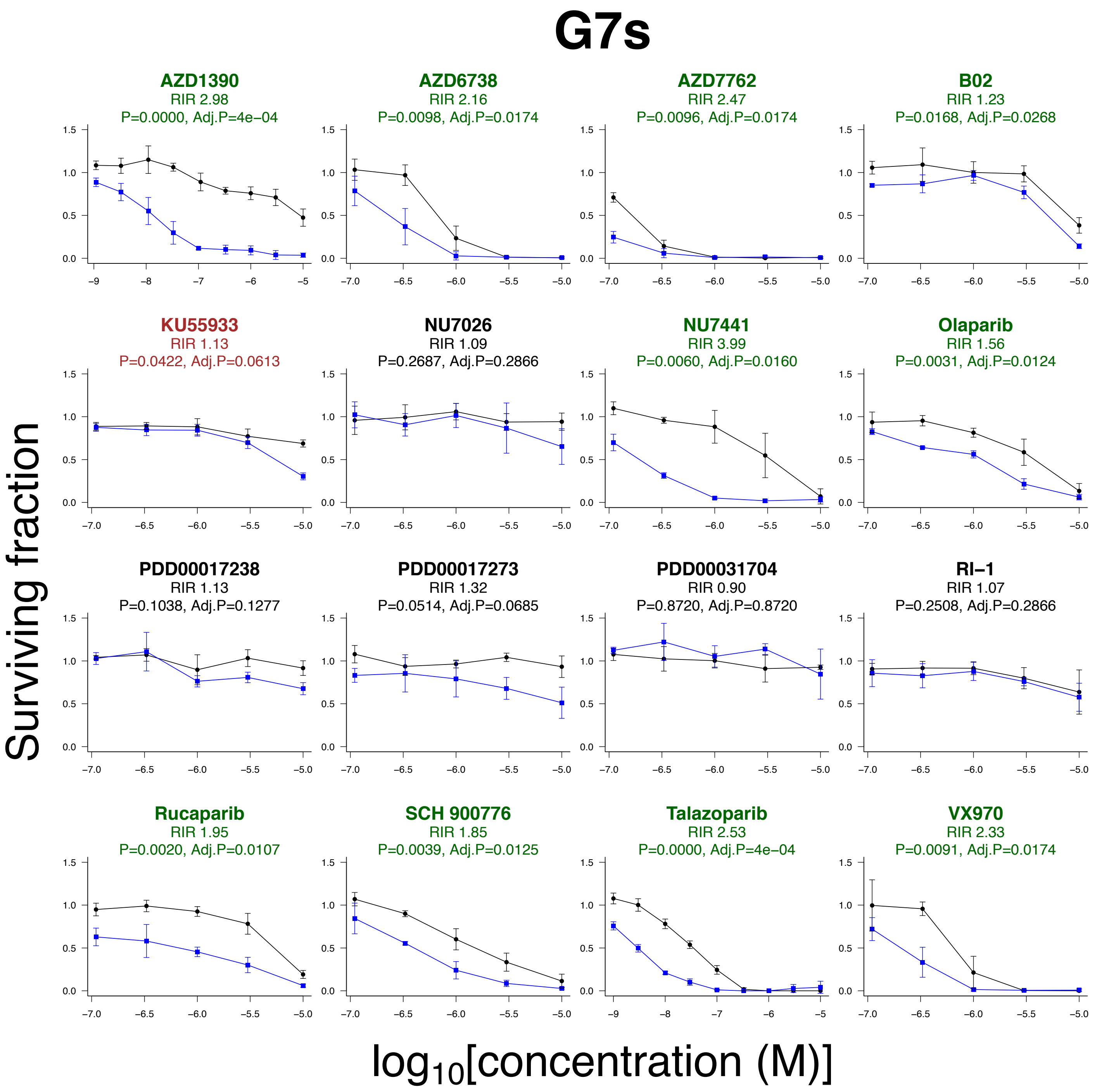


**Supp. Fig. S3: Dose response curves for DNA damage response inhibitors screened in G7s cells.**

A panel of DNA damage response (DDR) inhibitors was screened for interaction with radiation using the ClonoScreen3D platform in G7s cells. Cells were incubated with drugs for two hours prior to irradiation (3 Gy). Following normalization for the effect of IR alone, radiation interaction ratio (RIR) values were computed and tested using one-tailed ratio t-testing, with or without FDR P-value adjustment. *n*=3.

Supp. Figure S4

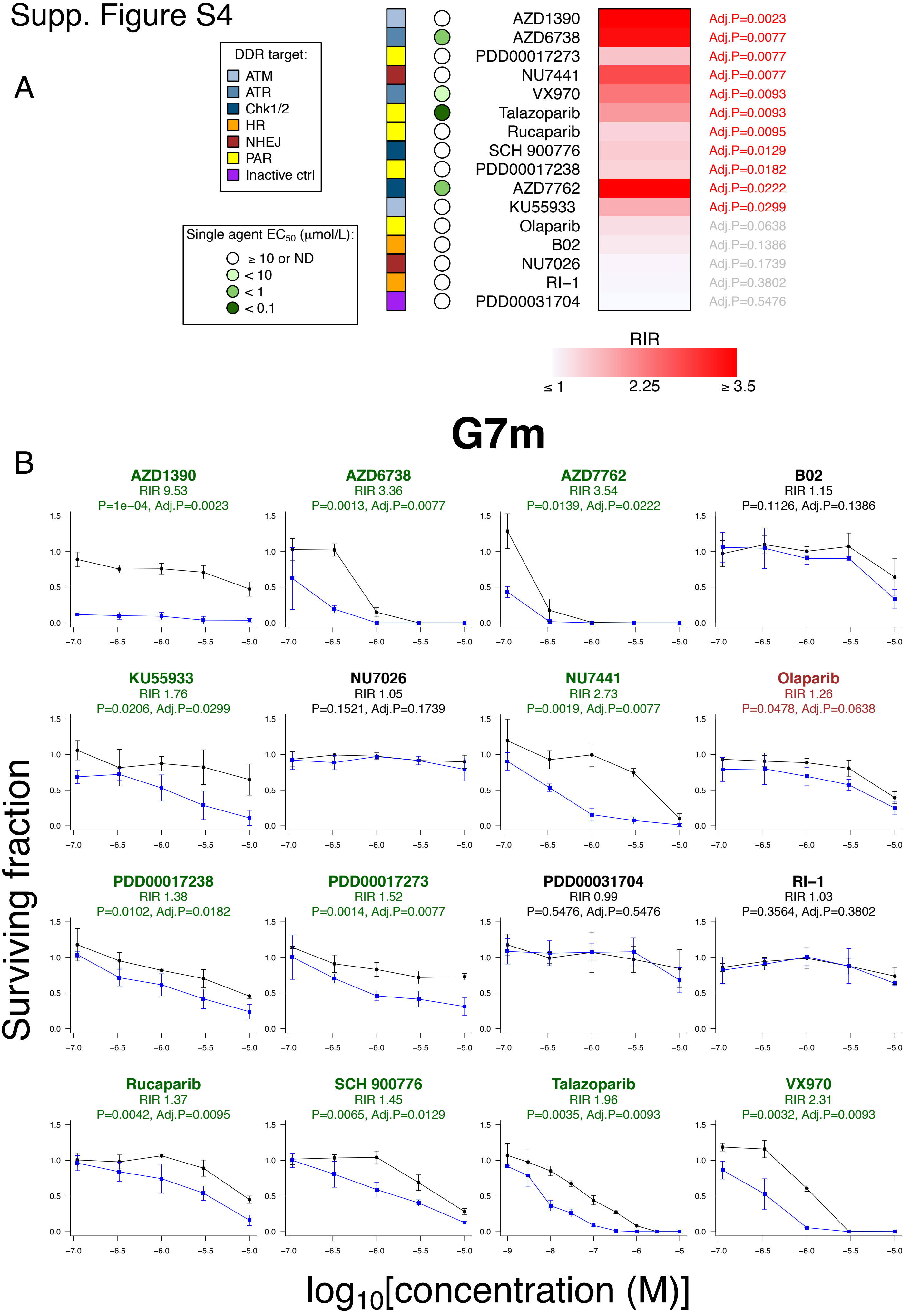

**Supp. Fig. S4: Quantitative comparison of the radiation interaction of DNA damage response inhibitors using the ClonoScreen3D platform in G7m cells.**

**A,** A panel of DNA damage response (DDR) inhibitors was screened for interaction with radiation using the ClonoScreen3D platform in G7m cells. Cells were incubated with drugs for two hours prior to irradiation (3 Gy). Following normalization for the effect of IR alone, radiation interaction ratio (RIR) values were computed and compounds ranked by FDR adjusted P-value, following one-tailed ratio t-testing. Drug single agent activity was quantified as $EC_{50}$ following fitting of a 4-parameter dose response model to sham-irradiated samples. The target pathway or protein is indicated. HR homologous recombination, NHEJ non-homologous end-joining, PAR poly(ADP-ribose), ND not determined. Data generated in three independent experiments. **B**, Dose response curves for DNA damage response inhibitors screened in G7m cells.

Supp. Figure S5

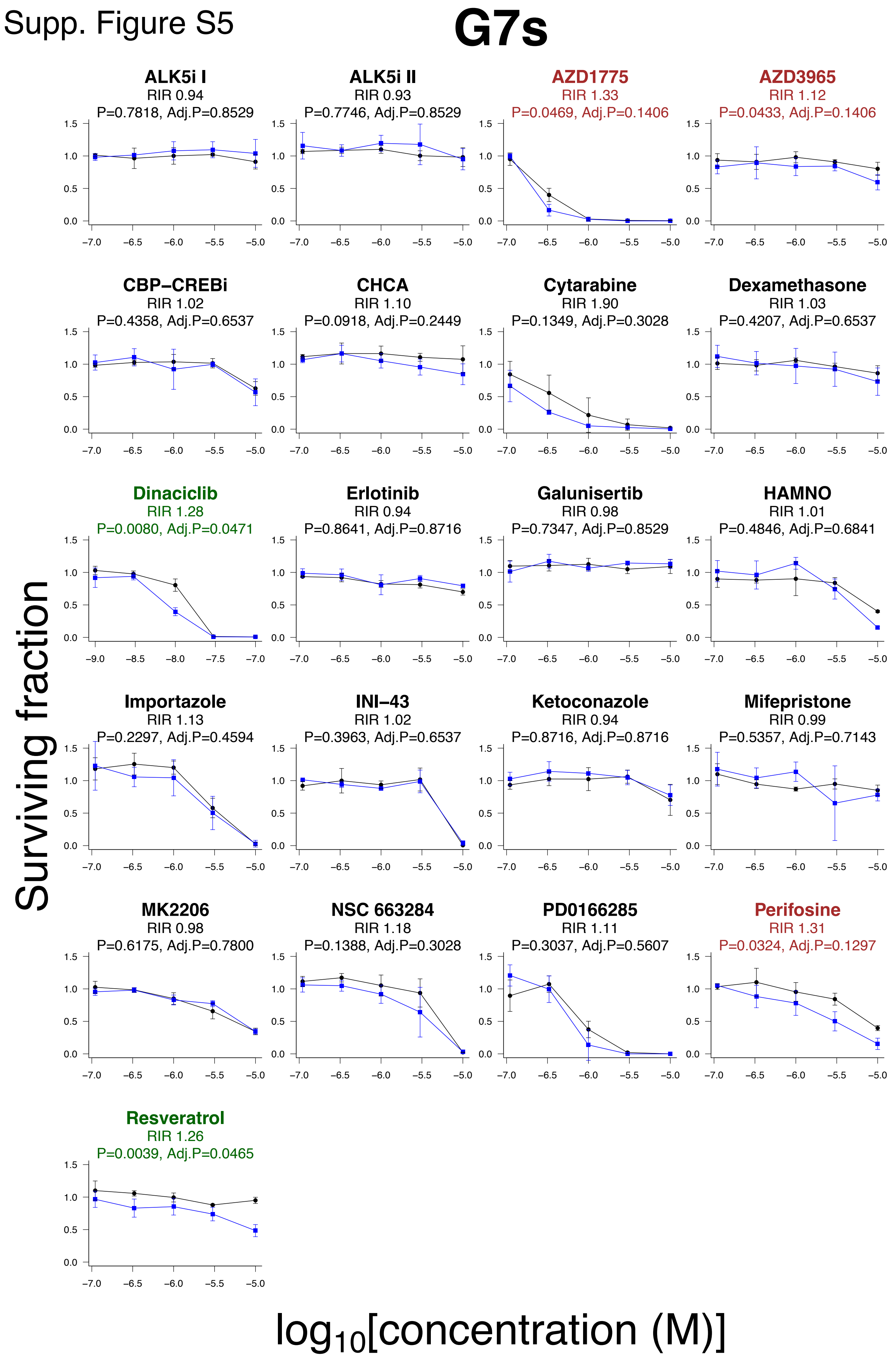

**Supp. Fig. S5: Dose response curves for compounds screened in G7s cells.**

A panel of commercially available inhibitors was screened for interaction with radiation using the ClonoScreen3D platform in G7s cells. Cells were incubated with drugs for two hours prior to irradiation (3 Gy). Following normalization for the effect of IR alone, radiation interaction ratio (RIR) values were computed and tested using one-tailed ratio t-testing, with or without FDR P-value adjustment. *n*=3.

Supp. Figure S6

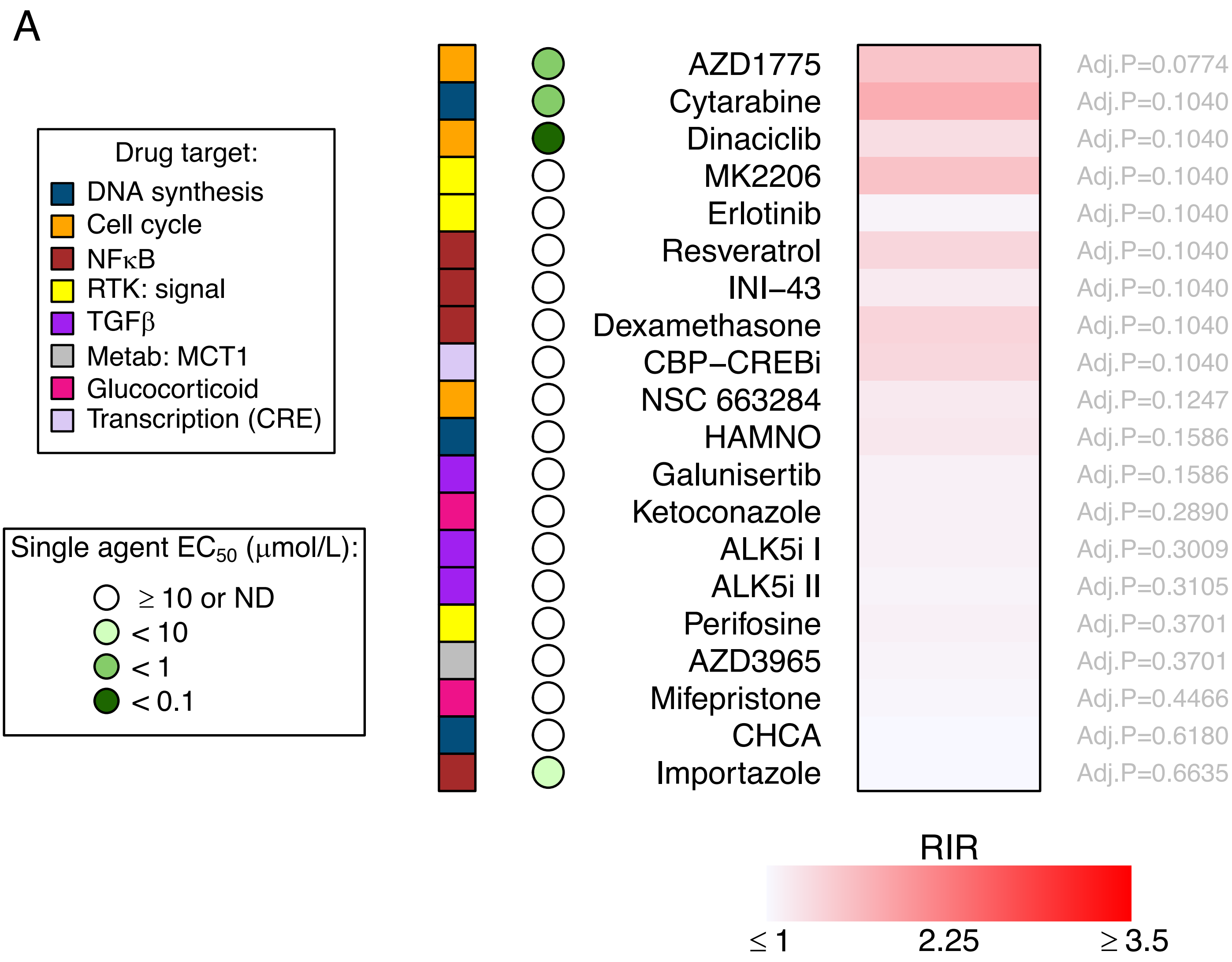

Supp. Figure S6

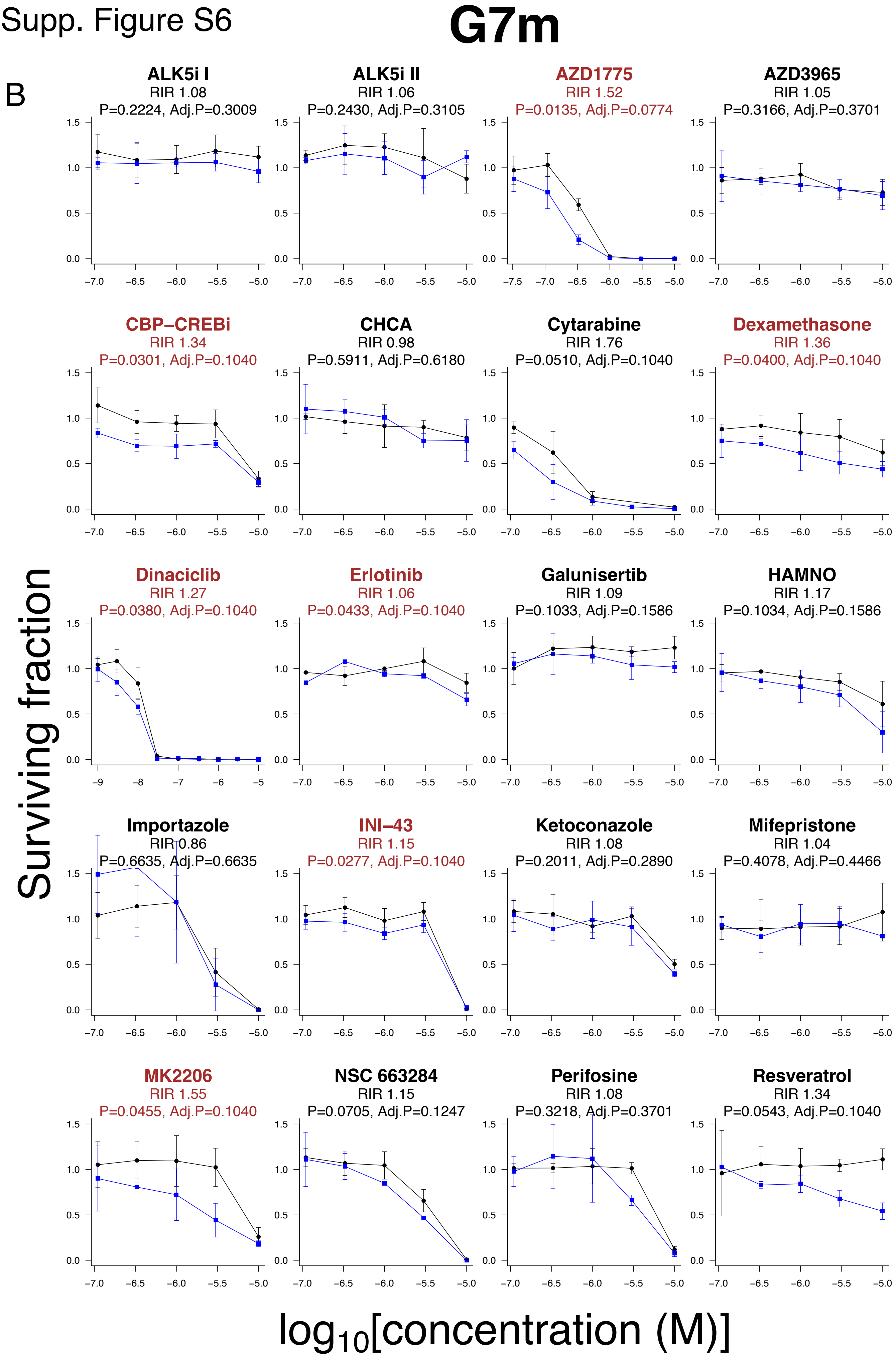

**Supp. Fig. S6: Evaluation of novel drugs-radiation combinations using the ClonoScreen3D platform for identification of novel radiosensitizers in G7m cells.**

**A**, A panel of commercially available inhibitors targeting pathways identified by transcriptomic analysis was screened for interaction with radiation using the ClonoScreen3D platform in G7s cells. Cells were incubated with drugs for two hours prior to irradiation (3 Gy). Following normalization for the effect of IR alone, radiation interaction ratio (RIR) values were computed and compounds ranked by FDR adjusted P-value, following one-tailed ratio t-testing. Drug single agent activity was quantified as $EC_{50}$ following fitting of a 4-parameter dose response model to sham-irradiated samples. The target pathway or protein is indicated. DDR DNA damage response, RTK receptor tyrosine kinase, Metab metabolism, CRE cAMP response element. Data generated in three independent experiments. **B**, Dose response curves for inhibitors screened in G7m cells.

### Supp. Figure S7

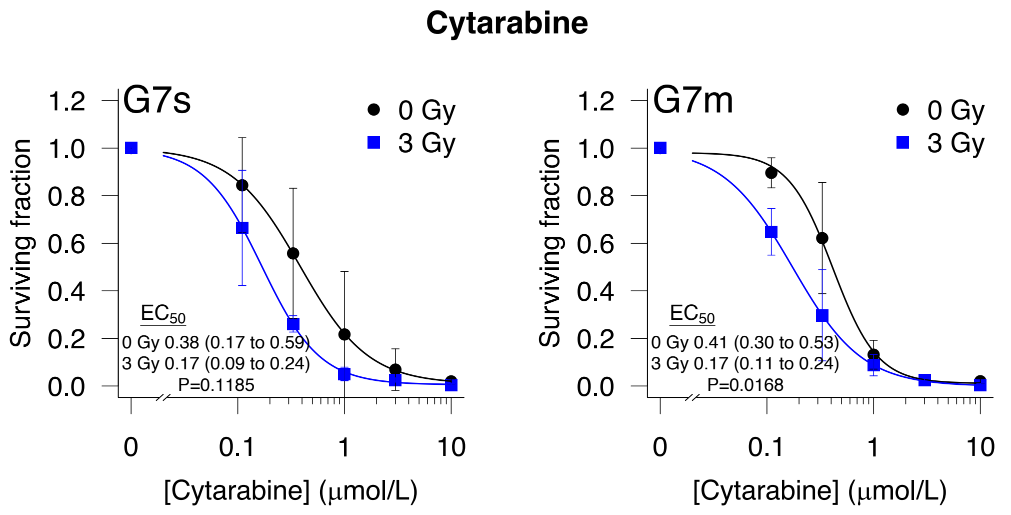


**Supp. Fig. S7: Additional comparison of dose response shift for cytarabine treated G7s and G7m cells.**
In addition to RIR analysis, data obtained in the ClonoScreen3D assay for G7s and G7m cells treated with cytarabine were fitted with a 4-parameter dose response curve, following normalization for the effect of IR. A shift in dose response was assessed by comparison of $EC_{50}$ between irradiated and sham-irradiated samples. EC50 (μmol/L) with 95% confidence interval calculated by fitting of a 4-parameter dose response curve, *n=3.*

#### Supplementary Table 1. Drug information.

| Drug name | CAS | Pathway | Main target gene | Other target gene | Other target gene | Other target gene | Other target gene | Other target gene | $IC_{50}$ (cell free) | $EC_{50}$ (cellular) | Supplier |
|---|---|---|---|---|---|---|---|---|---|---|---|
| ALK5i I | 396129-53-6 | TGFβ | TGFBR1 | | | | | | 51 nmole/L | | Enzo Life Sciences |
| ALK5i II | 446859-33-2 | TGFβ | TGFBR1 | | | | | | 23 nmole/L | | Enzo Life Sciences |
| AZD1775 | 955365-80-7 | Cell cycle | WEE1 | | | | | | 5.2 nmol/L | | Selleckchem |
| AZD3965 | 1448671-31-5 | Metabolism | MCT1 | MCT2 | | | | | | | Selleckchem |
| AZD6738 | 1352226-88-0 | DDR: signal | ATR | ATM | | | | | 1 nmole/L | | Selleckchem |
| AZD7762 | 860352-01-8 | DDR: signal | CHEK1 | CHEK2 | | | | | 5 nmole/L | | Selleckchem |
| B02 | 1290541-46-6 | DDR: repair HR | RAD51 | | | | | | 27.4 µmole/L | | Calbiochem/MERCK |
| CBP-CREBi | 92-78-4 | Transcription (CRE) | CBP | CREB | | | | | 2.9 µmole/L | | Calbiochem/MERCK |
| CHCA | 28166-41-8 | Metabolism | MCT1 | | | | | | 1.5 µmole/L | | Selleckchem |
| Cytarabine | 147-94-4 | DNA replication | | | | | | | | 16 nmole/L; CCRF-CEM cells | Selleckchem |
| Dexamethasone | 50-02-2 | NFkB | | | | | | | | 0.47 nmole/L; human PBMC | Selleckchem |
| Dinaciclib | 779353-01-4 | Cell cycle | CDK2 | CDK5 | CDK1 | CDK9 | | | 1 ; 1 ; 3 ; 4 nmole/L | | Selleckchem |
| Erlotinib | 183321-74-6 | RTK: signalling | EGFR | | | | | | 2 nmole/L | | Selleckchem |
| Galunisertib | 700874-72-2 | TGFβ | TGFBR1 | | | | | | 56 nmole/L | | Selleckchem |
| HAMNO | 138736-73-9 | DNA replication | RPA1 | | | | | | | | Selleckchem |
| Importazole | 662163-81-7 | NFκB | KPNB1 | NFkB | | | | | | 4.48 µmole/L; RPMI 8226 cells | Selleckchem |
| INI-43 | 881046-01-1 | NFκB | KPNB1 | NFY | AP-1 | p65 | NFAT | | | | Sigma-Aldrich/MERCK |
| Ketoconazole | 65277-42-1 | Glucocorticoid | CYP17A1 | CYP11B1 | | | | | 2.38 ; 0.608 µmole/L | | Sigma-Aldrich/MERCK |
| KU55933 | 587871-26-9 | DDR: signal | ATM | | | | | | 12.9 nmole/L | | Selleckchem |
| Methotrexate | 59-05-2 | DDR: MMR | DHFR | NF-kB | | | | | | 16 nmole/L; D54 cells | Selleckchem |
| Mifepristone | 84371-65-3 | Glucocorticoid | NR3C3 | NR3C1 | NR3C4 | | | | 0.025 ; 2.2 ; 10 nmole/L | | Sigma-Aldrich/MERCK |
| MK2206 | 1032350-13-2 | RTK: signalling | AKT1 | AKT2 | AKT3 | | | | 8 ; 12 ; 65 nmole/L | | Selleckchem |
| NSC 663284 | 383907-43-5 | Cell cycle | CDC25A | CDC25B | CDC25C | | | | NA ; 210 nmol/L; NA | | Sigma-Aldrich/MERCK |
| NU7026 | 154447-35-5 | DDR: repair NHEJ | PRKDC | PIK3CA | PIK3CD | PIK3CG | | | 230 nmole/L | | Selleckchem |
| NU7441 | 503468-95-9 | DDR: repair NHEJ | PRKDC | mTOR | PI3K | | | | 0.014; 1.7; 5 µmole/L | | Selleckchem |
| Olaparib | 763113-22-0 | DDR: repair BER | PARP1 | PARP2 | | | | | 5 ; 1 nmole/L | | Selleckchem |
| PD0166285 | 185039-89-8 | Cell cycle | WEE1 | MYT1 | CHEK1 | | | | 0.024 ; 0.072; 3.4 µmole/L | | Selleckchem |
| PDD00017238 | 1952247-05-0 | DDR: signal | PARG | | | | | | 40 nmole/L | | Ximbio |
| PDD00017273 | 1945950-21-9 | DDR: signal | PARG | | | | | | 26 nmole/L | | Ximbio |
| PDD00031704 | see ref. S1 | DDR: signal | PARG (inactive control) | | | | | | >100 µmole/L | | Ximbio |
| Perifosine | 157716-52-4 | RTK: signalling | AKT1 | AKT2 | AKT3 | | | | | 4.7 µmole/L; MM.1S cells | Selleckchem |
| Resveratrol | 501-36-0 | NFκB | SIRT1 | SIRT2 | NQO1 | IKKB | COX1 | COX2 | NA ; NA ; 0.088; 1; 1.1; 1.1 µmole/L | | Selleckchem |
| RI-1 | 415713-60-9 | DDR: repair HR | RAD51 | | | | | | 5 µmole/L | | Sigma-Aldrich/MERCK |
| Rucaparib | 283173-50-2 | DDR: PAR | PARP1 | | | | | | 1.4 nmole/L | | Selleckchem |
| SCH 900776 | 891494-63-6 | DDR: signal | CHEK1 | CHEK2 | | | | | 3; 160 nmole/L | | Selleckchem |
| SCR7 | 14892-97-8 | DDR: repair | LIG4 | | | | | | 200 µmole/L | 40 µmole/L; MCF7 cells | Selleckchem |
| Talazoparib | 1207456-01-6 | DDR: PAR | PARP1 | PARP2 | | | | | 0.57 nmole/L | | Selleckchem |
| VX970 | 1232416-25-9 | DDR: signal | ATR | ATM | | | | | 0.2 ; 34 nmole/L | | Selleckchem |

Reference S1. James, D.I., et al., First-in-Class Chemical Probes against Poly(ADP-ribose) Glycohydrolase (PARG) Inhibit DNA Repair with Differential Pharmacology to Olaparib. ACS Chem Biol, 2016. 11(11): p. 3179-3190.

**Supplementary Table 2. Sensitizer enhancement ratios elicited by PARP inhibitors.**

Full 3D radiation dose response clonogenic survival of G7s and G7m cells treated with olaparib (1 µmol/L), rucaparib (1 µmol/L) and talazoparib (5 nmol/L) two hours prior to IR. Data fitted using the linear quadratic model. Sensitizer enhancement ratios (SER) calculated using linear quadratic mean inactivation dose (MID) and subject to one-tailed ratio t-test. SER 95% confidence intervals shown.

| Cell | Drug | MID.veh | MID.drug | SER | SER. lower | SER. upper | P.val |
|---|---|---|---|---|---|---|---|
| G7s | Olaparib (1µmol/L) | 3.1199 | 2.5059 | 1.2450 | 1.0484 | 1.4560 | 0.0167 |
| G7s | Rucaparib (1µmol/L) | 3.1199 | 2.4368 | 1.2803 | 1.0355 | 1.6079 | 0.0141 |
| G7s | Talazoparib (5nmol/L) | 3.1199 | 1.8477 | 1.6886 | 1.0485 | 3.7976 | 0.0129 |
| G7m | Olaparib (1µmol/L) | 3.3912 | 2.7783 | 1.2206 | 0.9873 | 1.5951 | 0.0279 |
| G7m | Rucaparib (1µmol/L) | 3.3912 | 2.6459 | 1.2817 | 1.0666 | 1.6024 | 0.0124 |
| G7m | Talazoparib (5nmol/L) | 3.3912 | 1.9322 | 1.7551 | 1.1233 | 3.9973 | 0.0140 |

**Supplementary Table 3. Summary of ClonoScreen3D results for DNA damage response inhibitors in G7s cells.**

A panel of DNA damage response (DDR) inhibitors was screened for interaction with radiation using the ClonoScreen3D platform in G7s cells. Cells were incubated with drugs for two hours prior to irradiation (3 Gy). Radiation interaction ratio (RIR) values and FDR adjusted P-values were computed, following one-tailed ratio t-testing. Drug single agent activity was quantified as $EC_{50}$ (μmol/L) following fitting of a 4-parameter dose response model to sham-irradiated samples. The target pathway or protein is indicated. HR homologous recombination, NHEJ non-homologous end-joining, PAR poly(ADP-ribose), AUC area-under-the-curve. Data generated in three independent experiments. RIR 95% confidence intervals shown.

| Drug | Pathway | EC50.0Gy | EC50.0Gy .lower | EC50.0Gy .upper | AUC.0Gy | AUC.3Gy | RIR | RIR .lower | RIR .upper | P.val | Adj.P.val |
|---|---|---|---|---|---|---|---|---|---|---|---|
| AZD1390 | ATM | | | | 3.5685 | 1.1963 | 2.9829 | 2.2634 | 4.2299 | 0.0000 | 0.0004 |
| KU55933 | ATM | | | | 1.6280 | 1.4457 | 1.1261 | 0.9593 | 1.2934 | 0.0422 | 0.0613 |
| AZD6738 | ATR | 0.7240 | 0.5219 | 0.9261 | 0.8304 | 0.3846 | 2.1592 | 1.0645 | 17.2438 | 0.0098 | 0.0174 |
| VX970 | ATR | 0.7217 | 0.4645 | 0.9788 | 0.8001 | 0.3438 | 2.3275 | 1.2847 | 12.0652 | 0.0091 | 0.0174 |
| AZD7762 | Chk1/2 | 0.1572 | 0.1432 | 0.1712 | 0.2474 | 0.1002 | 2.4695 | | | 0.0096 | 0.0174 |
| SCH 900776 | Chk1/2 | 1.2415 | 0.6662 | 1.8167 | 1.1716 | 0.6328 | 1.8514 | 1.5303 | 2.1989 | 0.0039 | 0.0125 |
| B02 | HR | | | | 1.8478 | 1.5033 | 1.2292 | 1.0406 | 1.4405 | 0.0168 | 0.0268 |
| RI-1 | HR | | | | 1.6597 | 1.5518 | 1.0695 | 0.8175 | 1.3729 | 0.2508 | 0.2866 |
| PDD00031704 | Inactive ctrl | | | | 1.9251 | 2.1483 | 0.8961 | 0.7085 | 1.1294 | 0.8720 | 0.8720 |
| NU7026 | NHEJ | | | | 1.9276 | 1.7684 | 1.0900 | 0.7519 | 1.7431 | 0.2687 | 0.2866 |
| NU7441 | NHEJ | | | | 1.4369 | 0.3603 | 3.9876 | 2.5668 | 5.4087 | 0.0060 | 0.0160 |
| Olaparib | PAR | | | | 1.3981 | 0.8970 | 1.5586 | 1.3360 | 1.8025 | 0.0031 | 0.0124 |
| PDD00017238 | PAR | | | | 1.9469 | 1.7222 | 1.1305 | 0.8968 | 1.4504 | 0.1038 | 0.1277 |
| PDD00017273 | PAR | | | | 1.9329 | 1.4605 | 1.3234 | 0.8839 | 2.4897 | 0.0514 | 0.0685 |
| Rucaparib | PAR | | | | 1.5852 | 0.8129 | 1.9501 | 1.3634 | 3.2905 | 0.0020 | 0.0107 |
| Talazoparib | PAR | 0.0316 | 0.0265 | 0.0367 | 1.5534 | 0.6151 | 2.5252 | 2.1026 | 3.1091 | 0.0000 | 0.0004 |

**Supplementary Table 4. Summary of ClonoScreen3D results for DNA damage response inhibitors in G7m cells.**

A panel of DNA damage response (DDR) inhibitors was screened for interaction with radiation using the ClonoScreen3D platform in G7m cells. Cells were incubated with drugs for two hours prior to irradiation (3 Gy). Radiation interaction ratio (RIR) values and FDR adjusted P-values were computed, following one-tailed ratio t-testing. Drug single agent activity was quantified as $EC_{50}$ (μmol/L) following fitting of a 4-parameter dose response model to sham-irradiated samples. The target pathway or protein is indicated. HR homologous recombination, NHEJ non-homologous end-joining, PAR poly(ADP-ribose), AUC area-under-the-curve. Data generated in three independent experiments. RIR 95% confidence intervals shown.

| Drug | Pathway | EC50.0Gy | EC50.0Gy .lower | EC50.0Gy .upper | AUC.0Gy | AUC.3Gy | RIR | RIR .lower | RIR .upper | P.val | Adj.P.val |
|---|---|---|---|---|---|---|---|---|---|---|---|
| AZD1390 | ATM | | | | 1.4158 | 0.1485 | 9.5342 | | | 0.0001 | 0.0023 |
| KU55933 | ATM | | | | 1.6420 | 0.9345 | 1.7571 | 1.0447 | 3.2135 | 0.0206 | 0.0299 |
| AZD6738 | ATR | 0.8589 | -0.7037 | 2.4216 | 0.8059 | 0.2402 | 3.3553 | 1.6889 | 55.2394 | 0.0013 | 0.0077 |
| VX970 | ATR | 1.0092 | -0.8014 | 2.8199 | 1.1314 | 0.4905 | 2.3068 | 1.4080 | 5.9937 | 0.0032 | 0.0093 |
| AZD7762 | Chk1/2 | 0.2902 | -0.8039 | 1.3842 | 0.3942 | 0.1113 | 3.5428 | 1.9164 | 6.8417 | 0.0139 | 0.0222 |
| SCH 900776 | Chk1/2 | | | | 1.6543 | 1.1410 | 1.4498 | 1.1122 | 2.0107 | 0.0065 | 0.0129 |
| B02 | HR | | | | 1.9423 | 1.6821 | 1.1547 | 0.8555 | 1.6744 | 0.1126 | 0.1386 |
| RI-1 | HR | | | | 1.7636 | 1.7180 | 1.0266 | 0.8531 | 1.2445 | 0.3564 | 0.3802 |
| PDD00031704 | Inactive ctrl | | | | 1.9763 | 1.9977 | 0.9893 | 0.7803 | 1.2554 | 0.5476 | 0.5476 |
| NU7026 | NHEJ | | | | 1.8590 | 1.7741 | 1.0478 | 0.9308 | 1.1897 | 0.1521 | 0.1739 |
| NU7441 | NHEJ | | | | 1.6046 | 0.5869 | 2.7343 | 2.0912 | 3.6182 | 0.0019 | 0.0077 |
| Olaparib | PAR | | | | 1.5879 | 1.2556 | 1.2646 | 0.9115 | 1.9676 | 0.0478 | 0.0638 |
| PDD00017238 | PAR | | | | 1.6017 | 1.1584 | 1.3827 | 1.0663 | 1.8536 | 0.0102 | 0.0182 |
| PDD00017273 | PAR | | | | 1.6567 | 1.0869 | 1.5243 | 1.2426 | 1.9234 | 0.0014 | 0.0077 |
| Rucaparib | PAR | | | | 1.7818 | 1.2996 | 1.3710 | 1.1266 | 1.7084 | 0.0042 | 0.0095 |
| Talazoparib | PAR | 0.0761 | 0.0419 | 0.1103 | 1.9103 | 0.9760 | 1.9573 | 1.5783 | 2.3996 | 0.0035 | 0.0093 |

**Supplementary Table 5. Summary of ClonoScreen3D results for commercially available inhibitors in G7s cells.**

A panel of commercially available inhibitors was screened for interaction with radiation using the ClonoScreen3D platform in G7s cells. Cells were incubated with drugs for two hours prior to irradiation (3 Gy). Radiation interaction ratio (RIR) values and FDR adjusted P-values were computed, following one-tailed ratio t-testing. Drug single agent activity was quantified as $EC_{50}$ (μmol/L) following fitting of a 4-parameter dose response model to sham-irradiated samples. The target pathway or protein is indicated. DDR DNA damage response, Metab metabolism, RTK receptor tyrosine kinase, CRE cAMP response element. Data generated in three independent experiments. RIR 95% confidence intervals shown.

| Drug | Pathway | EC50.0Gy | EC50.0Gy .lower | EC50.0Gy .upper | AUC.0Gy | AUC.3Gy | RIR | RIR .lower | RIR .upper | P.val | Adj.P.val |
|---|---|---|---|---|---|---|---|---|---|---|---|
| AZD1775 | Cell cycle | 0.2873 | 0.2715 | 0.3030 | 0.4374 | 0.3289 | 1.3299 | 0.8991 | 2.4733 | 0.0469 | 0.1406 |
| Dinaciclib | Cell cycle | 0.0128 | 0.0044 | 0.0213 | 1.1454 | 0.8922 | 1.2838 | 1.0852 | 1.5209 | 0.0080 | 0.0471 |
| NSC 663284 | Cell cycle | | | | 1.8066 | 1.5250 | 1.1846 | 0.8024 | 1.9003 | 0.1388 | 0.3028 |
| PD0166285 | Cell cycle | 0.9476 | 0.3307 | 1.5645 | 0.9182 | 0.8301 | 1.1062 | 0.6359 | 3.6671 | 0.3037 | 0.5607 |
| Cytarabine | DNA synthesis | 0.3833 | 0.3549 | 0.4118 | 0.6118 | 0.3215 | 1.9028 | -0.0657 | 5.0960 | 0.1349 | 0.3028 |
| HAMNO | DNA synthesis | | | | 1.5933 | 1.5832 | 1.0064 | 0.6539 | 1.7341 | 0.4846 | 0.6841 |
| Ketoconazole | Glucocorticoid | | | | 1.9197 | 2.0521 | 0.9355 | 0.8049 | 1.0795 | 0.8716 | 0.8716 |
| Mifepristone | Glucocorticoid | | | | 1.8295 | 1.8555 | 0.9860 | 0.6281 | 2.1503 | 0.5357 | 0.7143 |
| AZD3965 | Metab: MCT1 | | | | 1.7926 | 1.6052 | 1.1168 | 0.9708 | 1.2960 | 0.0433 | 0.1406 |
| CHCA | Metab: MCT1 | | | | 2.2158 | 2.0154 | 1.0994 | 0.9110 | 1.2964 | 0.0918 | 0.2449 |
| Dexamethasone | NFκB | | | | 1.9266 | 1.8723 | 1.0290 | 0.6834 | 1.9165 | 0.4207 | 0.6537 |
| Importazole | NFκB | | | | 1.7548 | 1.5584 | 1.1260 | 0.7056 | 2.7757 | 0.2297 | 0.4594 |
| INI-43 | NFκB | | | | 1.6569 | 1.6189 | 1.0235 | 0.7803 | 1.2974 | 0.3963 | 0.6537 |
| Resveratrol | NFκB | | | | 1.9337 | 1.5334 | 1.2611 | 1.1012 | 1.4530 | 0.0039 | 0.0465 |
| Erlotinib | RTK: signal | | | | 1.6462 | 1.7461 | 0.9428 | 0.8255 | 1.0900 | 0.8641 | 0.8716 |
| MK2206 | RTK: signal | | | | 1.5424 | 1.5713 | 0.9816 | 0.7971 | 1.1770 | 0.6175 | 0.7800 |
| Perifosine | RTK: signal | | | | 1.7561 | 1.3388 | 1.3117 | 0.9578 | 1.8824 | 0.0324 | 0.1297 |
| ALK5i I | TGFβ | | | | 1.9315 | 2.0567 | 0.9391 | 0.7666 | 1.1511 | 0.7818 | 0.8529 |
| ALK5i II | TGFβ | | | | 2.0618 | 2.2054 | 0.9349 | 0.7362 | 1.2522 | 0.7746 | 0.8529 |
| Galunisertib | TGFβ | | | | 2.1412 | 2.1824 | 0.9811 | 0.9050 | 1.0616 | 0.7347 | 0.8529 |
| CBP-CREBi | Transcription (CRE) | | | | 1.8947 | 1.8633 | 1.0168 | 0.7499 | 1.5237 | 0.4358 | 0.6537 |

**Supplementary Table 6. Summary of ClonoScreen3D results for commercially available inhibitors in G7m cells.**

A panel of commercially available inhibitors was screened for interaction with radiation using the ClonoScreen3D platform in G7m cells. Cells were incubated with drugs for two hours prior to irradiation (3 Gy). Radiation interaction ratio (RIR) values and FDR adjusted P-values were computed, following one-tailed ratio t-testing. Drug single agent activity was quantified as $EC_{50}$ (µmol/L) following fitting of a 4-parameter dose response model to sham-irradiated samples. The target pathway or protein is indicated. DDR DNA damage response, Metab metabolism, RTK receptor tyrosine kinase, CRE cAMP response element. Data generated in three independent experiments. RIR 95% confidence intervals shown.

| Drug | Pathway | EC50.0Gy | EC50.0Gy .lower | EC50.0Gy .upper | AUC.0Gy | AUC.3Gy | RIR | RIR .lower | RIR .upper | P.val | Adj.P.val |
|---|---|---|---|---|---|---|---|---|---|---|---|
| AZD1775 | Cell cycle | 0.3461 | -0.1750 | 0.8672 | 1.0643 | 0.6984 | 1.5240 | 1.0405 | 2.6402 | 0.0135 | 0.0774 |
| Dinaciclib | Cell cycle | 0.0137 | 0.0108 | 0.0167 | 1.2327 | 0.9680 | 1.2736 | 0.9719 | 1.6224 | 0.0380 | 0.1040 |
| NSC 663284 | Cell cycle | | | | 1.6125 | 1.3967 | 1.1545 | 0.8851 | 1.4283 | 0.0705 | 0.1247 |
| CHCA | DNA synthesis | | | | 1.7957 | 1.8327 | 0.9798 | 0.7601 | 1.3399 | 0.5911 | 0.6180 |
| Cytarabine | DNA synthesis | 0.4143 | 0.3408 | 0.4879 | 0.6191 | 0.3522 | 1.7580 | 0.8058 | 5.1563 | 0.0510 | 0.1040 |
| HAMNO | DNA synthesis | | | | 1.7100 | 1.4593 | 1.1717 | 0.8625 | 1.6797 | 0.1034 | 0.1586 |
| Ketoconazole | Glucocorticoid | | | | 1.8503 | 1.7099 | 1.0821 | 0.8349 | 1.3643 | 0.2011 | 0.2890 |
| Mifepristone | Glucocorticoid | | | | 1.8189 | 1.7512 | 1.0387 | 0.4757 | 1.6382 | 0.4078 | 0.4466 |
| AZD3965 | Metab: MCT1 | | | | 1.6399 | 1.5616 | 1.0501 | 0.8025 | 1.4186 | 0.3166 | 0.3701 |
| Dexamethasone | NFκB | | | | 1.6121 | 1.1838 | 1.3618 | 0.9399 | 2.0236 | 0.0400 | 0.1040 |
| Importazole | NFκB | 2.8251 | -0.1539 | 5.8042 | 1.5702 | 1.8156 | 0.8648 | | | 0.6635 | 0.6635 |
| INI-43 | NFκB | | | | 1.8023 | 1.5716 | 1.1467 | 0.9889 | 1.3577 | 0.0277 | 0.1040 |
| Resveratrol | NFκB | | | | 2.0443 | 1.5247 | 1.3408 | 0.8591 | 1.8529 | 0.0543 | 0.1040 |
| Erlotinib | RTK: signal | | | | 1.9091 | 1.8025 | 1.0591 | 0.9791 | 1.1392 | 0.0433 | 0.1040 |
| MK2206 | RTK: signal | | | | 1.8806 | 1.2154 | 1.5473 | 0.8865 | 2.9326 | 0.0455 | 0.1040 |
| Perifosine | RTK: signal | | | | 1.7611 | 1.6251 | 1.0836 | 0.6563 | 2.8857 | 0.3218 | 0.3701 |
| ALK5i I | TGFβ | | | | 2.2065 | 2.0379 | 1.0827 | 0.7633 | 1.4246 | 0.2224 | 0.3009 |
| ALK5i II | TGFβ | | | | 2.2400 | 2.1227 | 1.0552 | 0.8678 | 1.2912 | 0.2430 | 0.3105 |
| Galunisertib | TGFβ | | | | 2.3280 | 2.1398 | 1.0879 | 0.9218 | 1.2677 | 0.1033 | 0.1586 |
| CBP-CREBi | Transcription (CRE) | | | | 1.7382 | 1.2993 | 1.3378 | 1.0041 | 1.7189 | 0.0301 | 0.1040 |

**Supplementary Table 7. Sensitizer enhancement ratios elicited by dinaciclib and cytarabine.**

Full 3D radiation dose response clonogenic survival of G7s, G7m and E2 cells treated with dinaciclib (1 and 10 nmol/L), and cytarabine (50 and 100 nmol/L) two hours prior to IR. Data fitted using the linear quadratic model. Sensitizer enhancement ratios (SER) calculated using linear quadratic mean inactivation dose (MID) and subject to one-tailed ratio t-test. SER 95% confidence intervals shown.

| Cell | Drug | MID.veh | MID.drug | SER | SER. lower | SER. upper | P.val |
|---|---|---|---|---|---|---|---|
| G7s | Dinaciclib (1 nmol/L) | 3.3368 | 3.2490 | 1.0270 | 0.8150 | 1.2653 | 0.3673 |
| G7s | Dinaciclib (10 nmol/L) | 3.3368 | 2.2706 | 1.4696 | 1.1690 | 1.8181 | 0.0100 |
| G7s | Cytarabine (50 nmol/L) | 3.3368 | 2.7798 | 1.2004 | 0.8141 | 1.9843 | 0.1188 |
| G7s | Cytarabine (100 nmol/L) | 3.3368 | 2.4709 | 1.3504 | 1.0616 | 1.6647 | 0.0197 |
| G7m | Dinaciclib (1 nmol/L) | 3.2484 | 3.6372 | 0.8931 | 0.8515 | 0.9365 | 0.9987 |
| G7m | Dinaciclib (10 nmol/L) | 3.2484 | 2.5975 | 1.2506 | 1.1334 | 1.3893 | 0.0018 |
| G7m | Cytarabine (50 nmol/L) | 3.2484 | 3.5253 | 0.9215 | NA | NA | 0.7139 |
| G7m | Cytarabine (100 nmol/L) | 3.2484 | 3.1479 | 1.0320 | 0.8072 | 1.3947 | 0.2523 |
| E2 | Dinaciclib (1 nmol/L) | 2.8384 | 2.4193 | 1.1732 | 0.9823 | 1.4156 | 0.0311 |
| E2 | Dinaciclib (10 nmol/L) | 2.8384 | 1.8321 | 1.5492 | 1.3406 | 1.7894 | 0.0013 |
| E2 | Cytarabine (50 nmol/L) | 2.8691 | 2.1312 | 1.3463 | 1.2074 | NA | 0.0006 |

**Supplementary Material 1. R script for computation of RIR from replicate clonogenic survival data.**

```
library(MESS)
library(mratios)


drug.conc <- c(0, 0.01, 0.03, 0.1, 0.3, 1, 3, 10)

sf.0gy <- data.frame(Rep1=c(1, 0.98, 0.92, 0.95, 0.83, 0.89, 0.94, 0.32),
                     Rep2=c(1, 0.96, 0.95, 0.89, 0.90, 0.83, 0.78, 0.40),
                     Rep3=c(1, 0.99, 0.91, 0.76, 0.86, 0.92, 0.84, 0.45))

sf.xgy <- data.frame(Rep1=c(1, 0.98, 0.92, 0.74, 0.23, 0.18, 0.14, 0.12),
                     Rep2=c(1, 0.96, 0.95, 0.59, 0.20, 0.23, 0.19, 0.16),
                     Rep3=c(1, 0.99, 0.91, 0.76, 0.26, 0.20, 0.17, 0.18))
## NB. `sf.xgy` data normalized for effect of radiation alone - SF calculated using plating efficiency of xGy 0mol/L sample(s).


RIR <- function(drug.conc, dat.0Gy, dat.xGy, logged.conc=F, rho=1, test.altHyp=c("greater", "two.sided")[1])
{
 if (!logged.conc & 0 %in% drug.conc)
  {
   stop("Remove 0 concentration")
  } else if (logged.conc) {
   log.x <- drug.conc
  } else {
   log.x <- log10(drug.conc)
  }
  auc.0gy <- apply(dat.0Gy, 2, auc, x=log.x)
  auc.xgy <- apply(dat.xGy, 2, auc, x=log.x)

  rir.res <- ttestratio(auc.0gy, auc.xgy, log=F, rho=rho, alternative=test.altHyp)
  return(rir.res)
```

```
}

## remove 0mol/L data

RIR(drug.conc=drug.conc[-1], dat.0Gy=sf.0gy[-1,], dat.xGy=sf.xgy[-1,], logged.conc=F)

##
##  Ratio t-test for unequal variances
##
## data:  x and y
## t = 31.573, df = 3.986, p-value = 3.101e-06
## alternative hypothesis: true ratio of means is greater than 1
## 95 percent confidence interval:
##  1.743013      Inf
## sample estimates:
##   mean x   mean y      x/y
## 2.529956 1.384748 1.827015

## RIR given as x/y estimate (0Gy/xGy)
```
